# Regulation of cyanobacterial type IV pilus-dependent functions by interaction between a c-di-GMP receptor and two transcription factors

**DOI:** 10.64898/2026.03.27.713163

**Authors:** Thomas Wallner, Chenliu He, Sherihan Samir, Eduardo Sabatine Lopes, Xiaoli Zeng, Cheng-Cai Zhang, Khaled A. Selim, Yiling Yang, Annegret Wilde

## Abstract

Cyanobacteria utilize type IV pili for many behavioural responses, such as phototaxis, aggregation, floating, and DNA uptake. Type IV pilus-dependent functions are regulated by the nucleotide second messengers, c-di-GMP and cAMP. In this study, we investigated the role of a recently identified c-di-GMP receptor (CdgR) in cyanobacteria that harbours a ComFB domain. ComFB-domain proteins are widespread in cyanobacteria and are also present in heterotrophic bacteria. We demonstrated that the CdgR homolog from the cyanobacterium *Synechocystis* sp. PCC 6803, a model organism for studying type IV pilus-dependent functions, specifically binds to c-di-GMP. Genetic and phenotypic analyses revealed that *Synechocystis* CdgR is involved in phototactic motility and natural competence. Inactivation of *cdgR* resulted in altered expression of specific sets of minor pilins, which are essential for motility or natural competence. We identified interactions between CdgR and the CRP-family transcription factors, SyCRP1 and SyCRP2. Disruption of these CdgR-SyCRP1 and CdgR/SyCRP2 complexes is initiated by elevated c-di-GMP levels. Moreover, the assembly and stability of these complexes are influenced by other cyclic nucleotides, such as cAMP and c-di-AMP. These observed interactions imply a complex regulatory mechanism by which CdgR influences gene expression in response to cyclic nucleotide messenger signalling, particularly c-di-GMP. The present findings highlight the importance of CdgR in c-di-GMP signalling and its role in regulating type IV pilus-dependent functions in *Synechocystis.* The modulation of the expression of specific minor pilin genes by CdgR, through interactions with the transcription factors SyCRP1 and SyCRP2, contributes to the establishment of multiple type IV pilus functions and adaptive behaviours of cyanobacteria.

## Introduction

Cyanobacteria, one of the most ancient organisms on Earth, are proficient in oxygenic photosynthesis and have colonized nearly all habitats. Many cyanobacterial species can move on surfaces using type IV pili (Bhaya et al., 2000; Chau et al., 2015). Motility confers adaptive advantages to cyanobacteria, enabling them to thrive in diverse and changing environments. Among these motility behaviors, phototaxis, which involves directional movement in response to light, has been extensively studied in *Synechocystis* sp. PCC 6803 (hereafter *Synechocystis*). In addition to motility, type IV pili play crucial roles in other cellular functions, including natural competence and cell-cell aggregation/flocculation (Conradi et al., 2019; Oeser et al., 2021). Disruption of *pilA1* results in the loss of type IV pili and type IV pilus-dependent functions (Bhaya et al., 2001). Besides the major pilin gene *pilA1*, the *Synechocystis* genome also contains multiple *pilA*-like genes encoding minor pilins (Linhartová et al., 2014). Major and minor pilins share a conserved cleavage site for maturation by the prepilin peptidase PilD and a similar secondary structure with a conserved N-terminal hydrophobic handle and a non-conserved C-terminal bulky head (Giltner et al., 2012). In *Synechocystis*, disruption of the gene cluster *pilA9*-*slr2019* leads to loss of flocculation and motility, while disruption of *pilA5* causes the loss of natural competence (Conradi et al., 2019; Wallner et al., 2020; Oeser et al., 2021). The expression of these minor pilins has been shown to be controlled by nucleotide second messengers (Savakis et al., 2012; Song et al., 2018; Wallner et al., 2020), however through yet unknown mechanism.

Similar to other bacteria, motility of *Synechocystis* depends on the second messenger cyclic dimeric guanosine monophosphate (c-di-GMP) (Savakis et al., 2012; Jenal et al., 2017; Mantovani et al., 2023). Intracellular c-di-GMP levels are balanced by two types of enzymes: diguanylate cyclases (DGC), which produce c-di-GMP, and phosphodiesterases (PDE), which degrade it (Römling et al., 2013). Elevated intracellular c-di-GMP levels are typically correlated with reduced motility and increased biofilm formation in numerous bacterial species (Hengge, 2009; Römling et al., 2013; Jenal et al., 2017; Hengge et al., 2023). Notably, in *Synechocystis*, blue light illumination mediated by the photoreceptor Cph2 inhibits twitching motility and induces cellular aggregation by increasing c-di-GMP levels, resulting in changes in gene expression (Savakis et al., 2012; Wallner et al., 2020). Although *Synechocystis* encodes potential c-di-GMP effector proteins, such as those with inactive GGDEF or MshEN domains (Chou and Galperin, 2016), none have been characterized yet. However, a c-di-GMP receptor, CdgR, has been described in the filamentous cyanobacterium *Anabaena* (*Nostoc*) sp. PCC 7120 (hereafter *Anabaena*) (Zeng et al., 2023). CdgR plays a critical role in regulating cell size in *Anabaena*, a filamentous cyanobacterium capable of differentiating into heterocyst for atmospheric nitrogen fixation under nitrogen-depleted conditions. In *Anabaena*, reduced levels of c-di-GMP or the accumulation of the apo-form of CdgR led to reduced cell size; however, when the c-di-GMP levels increased or CdgR was saturated by c-di-GMP, no effect on cell size was observed. The cell size control effect is mediated by CdgR through the interaction between CdgR and DevH, a CRP-family transcription factor that seemingly participates in the c-di-GMP signaling pathway in *Anabaena* via a mechanism that is not yet fully understood. CdgR is highly conserved in cyanobacteria, and the structure of its homolog in *Synechocystis* (Slr1970) in complex with c-di-GMP has been determined by crystallography (Zeng et al., 2023; Samir et al., 2025b). Furthermore, sequence and structural analyses suggested that CdgR has homologs in heterotrophic bacteria, such as *Bacillus subtilis* and *Vibrio cholerae* (Samir et al., 2025a) which belong to a widespread protein family harbouring a ComFB domain (Samir et al., 2025b). The functions of CdgR/ComFB homologs have been related to the control of motility and DNA uptake with the help of type IV pili (Hahn et al., 2025; Samir et al., 2025b), however the mechanism is still unknown.

The *Synechocystis* genome contains four genes encoding proteins with a ComFB-like domain: Slr1970, Sll1170, Sll1739, and Slr1505 (Samir et al., 2025b). All of these proteins combine the ComFB domain with additional N- or C-terminal extensions. To discriminate between these potential c-di-NMP receptor proteins, in the following, we use the initially introduced gene name *cdgR* for the *Synechocystis* homolog (Slr1970), which has the highest similarity to the *Anabaena* c-di-GMP receptor. Given the crucial role of c-di-GMP signaling in type IV pilus-dependent functions in *Synechocystis* (Enomoto et al., 2023; Mantovani et al., 2023), we investigated whether CdgR participates in the control of motility, aggregation, and natural competence in this model strain. We present evidence that *Synechocystis* CdgR is involved in controlling positive phototaxis and natural competence. Our investigations revealed that the absence of CdgR results in altered expression of minor pilin genes, and that this regulation is mediated by the interaction of CdgR with the CRP-like transcription factors SyCRP2 and SyCRP1. Based on our data, we propose a model for the control of minor pilin gene expression by c-di-GMP.

## Materials and methods

### Strains and growth conditions

All primers, plasmids, and strains used in this study are listed in Supplementary Tables S1-S3. The *Synechocystis* strain was cultured and maintained at 30 ℃ on BG11 agar or in liquid medium (Rippka et al., 1979). For the bacterial adenylate cyclase two-hybrid assay (BACTH), *E. coli* BTH101 (Euromedex) was utilized. It was grown and maintained at 37 ℃ on lysogeny broth (LB) agar or in LB broth (1% (w/v) tryptone, 0.5% (w/v) yeast extract, 0.5% (w/v) sodium chloride, pH 7). Unless otherwise stated, all LB plates contained 1.2% agar. When necessary, kanamycin or spectinomycin/streptomycin was added to the BG11 medium at final concentrations of 25 µg/mL and 5 µg/mL, respectively. Ampicillin, kanamycin, chloramphenicol, and spectinomycin were added to the LB medium at final concentrations of 100, 50, 25, and 50 µg/mL, respectively.

### Construction of plasmids and mutant strains

The *cdgR* (gene locus *slr1970*) was deactivated by replacing its entire open reading frame, except for the final 33 bp, with a kanamycin resistance cassette using advanced quick assembly cloning (Beyer et al., 2015). We amplified an upstream fragment containing the promoter of the transcriptional unit TU 1845 (Kopf et al., 2014) along with the RNA chaperone *hfq/ssr3341*, which is co-transcribed with *cdgR*. We used the primer pair AQ-*slr1970*-P1 and AQ-*slr1970*-P2 (Table S1) to amplify this fragment from *Synechocystis* wild-type chromosomal DNA. The downstream section, including the *slr1971* gene, was amplified from chromosomal DNA using the primer pair AQ-*slr1970*-P5/AQ-*slr1970*-P6. The kanamycin resistance cassette was amplified from the pUC4K plasmid (Vieira and Messing, 1982) using the primer pair AQ-*slr1970*-P3/AQ-*slr1970*-P4. The pUC19 plasmid (Yanisch-Perron et al., 1985) was used as the backbone for cloning assembly and was amplified using the primer pair AQ-*slr1970*-P7 and AQ-*slr1970*-P8. Following the protocol described by Ermakova-Gerdes and Vermaas (1999), the final construct was used to transform *Synechocystis* wild type. The resulting transformants were plated onto BG11 agar plates containing progressively higher concentrations of kanamycin, ranging from 1 μg/mL to 40 μg/mL. To detect complete segregation of the *cdgR::Km* mutant, colony PCR was performed using the primer pair, *slr1970*-col-fw and *slr1970*-col-rev.

To complement the *cdgR::Km* mutant, the entire transcriptional unit TU1845 (Kopf et al., 2014) was amplified from wild-type chromosomal DNA using the primer pair HindIII-1970-C-FW/BamHI-1970-C-REV. This process added BamHI and HindIII restriction enzyme recognition sequences to the 5’ and 3’ends of the DNA sequence, respectively. The amplified sequence was then subcloned into the pJET vector (Thermo Scientific). A NheI restriction enzyme recognition sequence was introduced in frame at alanine 169 position of CdgR using the Q5 site-directed mutagenesis kit (New England Biolabs) and primer pair A169A-NheI-fw and A169A-NheI-rev. A fused oligonucleotide (XbaI-FLAG-Fw and XbaI-FLAG-REV) encoding a triple FLAG tag, including a stop codon, was ligated into the open NheI site. The resulting construct was excised from the pJET vector using HindIII and BamHI and ligated into a modified conjugative pVZ322 plasmid. The resulting complementation plasmid, designated pEP-*cdgR*-FLAG, was conjugated into the *cdgR::Km* mutant using tri-parental mating.

The *ΔcdgR* in-frame deletion strain was generated using the established CRISPR-Cpf1-based genome editing (Niu et al., 2019). To construct an in-frame deletion of *cdgR*, a deletion plasmid, pCpf1b-*cdgR*-sp, was assembled. The left (LA) and right (RA) homology arms flanking the *cdgR* locus, along with the required guide sequence, were amplified from the template plasmid pCpf1-M*slr1970*R236. These fragments were ligated into the linearized pCpf1b-sp backbone. The resulting plasmid was transferred into wild-type cells via conjugation to generate the *ΔcdgR* strain. The *Δcph2* mutant was constructed via allelic exchange. The LA and RA fragments of *cph2* and the kanamycin resistance cassette (*aphI*) were assembled into a linearized pUC19 vector using a seamless cloning strategy, yielding the suicide plasmid pUC19-*cph2-km*. This plasmid was used for transformation of *Synechocystis*. To generate the Δ*cdgR*Δ*cph2* double mutant, the pUC19-*cph2-km* plasmid was transferred into the *ΔcdgR* background.

For genetic complementation and inducible overexpression, the open reading frames of *cdgR*, *ydjH, and yheH* were individually cloned into the pSCT3 vector. This shuttle vector contains a synthetic CT promoter, which is dually inducible by copper and theophylline. The resulting plasmids were designated pSCT3-*cdgR*, pSCT3-*ydeH* and pSCT3-*yhjH*. The complementation plasmid pSCT3-*cdgR* was introduced into the *ΔcdgR* strain to generate the complemented strain, C-Δ*cdgR*. For overexpression studies, respective plasmids were transferred into both wild type (pSCT3-*cdgR*, pSCT3-*ydeH* and pSCT3-*yhjH*) and Δ*cdgR* (pSCT3-*ydeH* and pSCT3-*yhjH*) backgrounds via conjugation. leading to the generation of WT_OE *cdgR*, WT_OE *yhjH,* WT_OE-*ydeH,* Δ*cdgR*_OE *yhjH* and Δ*cdgR*_OE *ydeH* strains.

### Phototaxis assay

Phototaxis assays were performed in accordance with the protocol previously outlined by Jakob et al. (2017). Wild-type and mutant strains were cultivated on solid BG11 plates (0.75-1.0% agar w/v) for 7-10 days before being resuspended in fresh BG11 to an OD_750_ of 15. In the standard phototaxis assay, BG11 containing 0.5-0.7% agar, 0.2% glucose, 10 mM Na_2_S_2_O_3_, and 12.5 mM TES was poured onto small square plates. For phototaxis assays performed on large square plates (24 cm x 24 cm), 0.3 µM CuSO_4_ and 2 mM theophylline were added to the medium to induce protein expression from the synthetic CT promoter. Subsequently, 2-µL aliquots of each cell culture were spotted onto the plate at regular intervals of 3 cm. To elicit phototactic movement, lateral light sources were provided using white, red (660 nm), green (520 nm) or blue LED (405 nm) strips. Phototactic responses were observed and documented after 5–12 days of incubation at 30 ℃.

### Analysis of single cell motility

Cells from the front of a moving colony were resuspended in fresh BG11 medium. Then, 3-µL aliquots were directly spotted on top of pre-warmed 5 mL BG11 motility medium solidified with 0.3 % (w/v) agarose and supplemented with 11 mM glucose and 10 mM TES (pH 8.9) in 35 mm glass-bottom plates (µDish, ibidi, Germany). The plates were maintained under dim light at 30 °C. After 5-10 min, a coverslip was precisely positioned on top of the cells. To reduce surface oscillations and prevent evaporation, the areas surrounding the plates were covered with a silicone ring. Time-lapse videos were captured at room temperature (ca. 22 °C) using an upright Nikon Eclipse Ni-U microscope (Nikon Instruments, Germany) fitted with a 40× objective lens (numerical aperture 0.75). Gradient illumination was performed using white light from the microscope condenser lamp. A red light-emitting diode (625 nm, World Trading Net GmbH, Germany) for directional illumination was mounted in a borehole of a black plastic cylinder surrounding the motility plate. Light intensity was measured using a LiCOR light meter with a planar quantum sensor (LI-COR Biosciences GmbH, Germany) with a detection window of 400–700 nm. The movement of individual cells was recorded at a rate of one frame per three seconds for three minutes. The exposure time was set to 200 ms to reduce the background light necessary for cell visualization.

### Single cell tracking and data analysis

Cell movement was tracked using Fiji software (Schindelin et al., 2012) and Fiji’s TrackMate plugin (Ershov et al., 2022) with the following parameter: the maximum distance of a detected cell between two frames was limited to 4 µm. It is imperative that the detected cells maintain their traceability for a minimum of three consecutive frames. Furthermore, the distance between cells must not exceed 4 µm in subsequent frames. The raw tracks were analyzed using R software and RStudio, with the CIRCULAR package implemented for data analysis (Agostinelli and Lund, 2023; R_Core_Team, 2023).

### Transmission electron microscopy

Wild-type and *ΔcdgR* mutant cells were cultivated at 30℃ until an OD_750_ of 0.6-1.0 was attained. Subsequently, 10 µL of cells were dispensed onto a parafilm layer, and a copper grid was carefully positioned on the surface of a droplet to allow cell attachment. After a 10-minute period, the grid with the attached cells was gently tapped with blotting paper at the edge to remove any liquid. Subsequently, the samples were transferred onto a droplet of 2% phosphotungstic acid solution (pH 7.0) for negative cell staining. Following a 3-minute incubation period, the copper grid was carefully removed and positioned on clean blotting paper. The air-dried specimens were examined at ambient temperature using a Hitachi transmission electron microscope operated at 80 kV.

### Flocculation assays

Flocculation assays were performed as described previously (Oeser et al., 2021). In summary, cultures were diluted to an OD_750nm_ of 0.25 in BG11 medium. Subsequently, 6 mL of each cell culture was added to a six-well plate (Corning Costar, non-treated, 392-0213, VWR, Germany) and incubated for 48 h at 30 °C and 40 μmol photons m^-2^ s^-1^ on an orbital shaker set at 95 rpm and an orbit of 10 mm (Multimix 1010, Heidolph Instruments, Germany). Images of the plates were captured before and after incubation using a Typhoon FLA4500 imaging system (GE Healthcare). The excitation of chlorophyll *a* was accomplished using laser light with a wavelength of 473 nm. The autofluorescence of *Synechocystis* cells was subsequently captured using a 665 nm longpass red emission filter. The photomultiplier tube was set to 500 V, and the pixel size was set to 25 μm. The aggregation score was calculated using the Quantity One Software (Bio-Rad Laboratories, Germany), as previously described (Conradi et al., 2019). A two-sample t-test assuming equal variances was applied.

### Transformation assays

Natural transformation competence was evaluated using a spectinomycin resistance-carrying suicide plasmid. 10 mL of a logarithmic culture (OD_750nm_ ∼0.85), were harvested (3,200 × *g*, RT, 10 min, cell pellets were resuspended in 300 µL of BG11, and 1 µg of plasmid DNA was added to the suspension. The cells were incubated for six hours at 30 °C under white light illumination (50 µmol photons m^-2^ s^-1^). Subsequently, 150 µL of the cells were streaked onto BG11 agar plates in duplicate. A small cellulose filter (2.5 cm diameter, Whatman qualitative filter, grade 1, 512-0375, Aventor) was positioned at the center of the plate. Then, 50 µL of a solution containing spectinomycin (50 mg/mL) was dispensed onto the filter to establish an antibiotic concentration gradient. The plates were incubated for 12 days, after which they were scanned using a flatbed scanner. The number of colony-forming units was determined and normalized to OD_750nm_ and the amount of DNA used in the assay. Integration of the transformation construct into the chromosome resulted in deletion of the RNA helicase gene *crhR/slr0083*. This deletion had no discernible effect under normal cultivation conditions (Prakash et al., 2010). A two-sample t-test assuming unequal variances was applied.

### RNA extraction

Cultures of *Synechocystis* wild-type and the *cdgR::Km* mutant strains were cultivated in BG11 medium and diluted to an OD_750nm_ of 0.35 in 50 mL BG11 medium. After 24 h of white-light exposure (70 µmol photons m^-2^ s^-1^) at 30 °C, cells were collected via filtration using polyethersulfone filter disks with 0.8 μm pore size (Pall, Germany). After filtration, the filters carrying the cells were instantly transferred into 1.6 mL PGTX solution (Pinto et al., 2009). RNA was isolated according to Pinto et al. (2009) with the following adjustments: the aqueous phase was extracted once more with 1-bromo-3-chloropropane following the initial extraction, and the phases were separated again. The RNA in the aqueous phase was precipitated at -20 °C using 2-propanol overnight. RNA was quantified using a NanoDrop ND2000 spectrophotometer (Thermo Fisher Scientific, Germany), and RNA integrity was assessed by separation on denaturing 1.3% (w/v) agarose/formaldehyde gels.

### Northern blot hybridization

Selected transcripts were verified by Northern blot hybridization using radioactively labeled RNA or DNA probes. The RNA probes were produced through in vitro transcription of PCR fragments using the Ambion T7 polymerase Maxiscript kit (Thermo Fisher Scientific, Germany) coupled with [α-^32^P]-UTP (Hartmann Analytics, Germany). The oligonucleotides used to amplify the respective probe DNA templates are listed in Table S1. A DNA probe for *Synechocystis* 16S rRNA (Table S1) was labeled with [α-^32^P]-dCTP (Hartmann Analytics, Germany) using the Rediprime II DNA labelling system (Cytiva, Germany). Total RNA obtained from individual experiments was isolated as described previously, followed by electrophoresis on 1.3% (w/v) agarose/formaldehyde gels and transfer onto Roti-Nylon plus membranes (Carl Roth, Germany) (Green and Sambrook, 2012). Hybridization with RNA or DNA probes was performed as described by Wallner et al. (2020). Signals were acquired using a Typhoon FLA4500 imaging system (GE Healthcare) and densitometrically analyzed using QuantityOne software (Bio-Rad).

### RNAseq analysis

The wild-type and Δ*cdgR* mutant strains were streaked onto BG11 plates and cultured under continuous light at an intensity of 30 μmol photons m^-2^ s^-1^ for one week. Subsequently, the cells were harvested from the plates, diluted 60-fold, and measured for OD_750_. Further dilution was carried out to achieve an OD of 15 using BG11 liquid medium, and aliquots were spotted onto the surface of BG11 agar medium (0.6% agar) in 10 cm square plates. The spots were placed in the middle row, approximately 7 cm from the bottom of the plate. Each plate was spotted with 50 µL of the diluted culture until completion, and three plates were prepared for each sample. The plates were then exposed to lateral white LED lamps with an intensity of approximately 50 μmol photons m^-2^ s^-1^ for 5 days. To collect samples for RNA-seq, plates were placed on ice and 1mL of pre-chilled PBS was added to wash off the cells, which were then collected into 1.5 mL tubes, with three tubes collected per plate. Following collection, the tubes were centrifuged at 6000 rpm for 3 min, and the supernatant was discarded. Cell pellets were rapidly frozen in liquid nitrogen and stored at -80 °C until further processing.

Total RNA was extracted, and ribosomal RNA was removed using the Ribo-clean rRNA Depletion Kit (Bacteria) (RN417, Vazyme, Nanjing, China). Subsequent sequencing libraries were constructed using the VAHTS Universal V8 RNA-seq Library Prep Kit for MGI (NRM605, Vazyme, Nanjing, China). The pooled libraries were sequenced on the MGI DNBSEQ-T7 platform at the Analytical and Testing Center of the Institute of Hydrobiology, Chinese Academy of Sciences. For RNA-seq analysis, raw sequencing data underwent quality filtering with Trimmomatic to remove low-quality reads and adapter sequences. Ribosomal RNA sequences were filtered out, resulting in clean data. These reads were subsequently aligned to the reference genome (GCF_000009725.1_ASM972v1 genomic.fna) using HISAT2 (version 2.2.1). StringTie was used to quantify the aligned reads. Finally, DESeq2 was used to analyze the count data and identify differentially expressed genes (DEGs), determining statistically significant changes in gene expression. The full data set is accessible in the Gene Expression Omnibus database with the accession number GSE322604 (https://www.ncbi.nlm.nih.gov/geo/query/acc.cgi?acc=GSE322604; security access token for reviewers: cxkpgyyavjshfyv)

### BACTH assay

Construction of the plasmids, cultivation of *E. coli* BTH101 and the experimental procedure of the BACTH asssy were performed as previously described (Karimova et al., 1998; Battesti and Bouveret, 2012; Selim et al., 2021) on LB-agar plates supplemented with X-gal (40 µg/mL), kanamycin (50 µg/mL), ampicillin (100 µg/mL) and IPTG (1 mM) or MacConkey agar plates supplemented with kanamycin (50 µg/mL), ampicillin (100 µg/mL) and IPTG (1 mM). The cells were spotted onto indicator plates and incubated at various temperatures (20, 25, 30, and 37 °C) for up to four days. The interaction between *Synechocystis* carbon control protein SbtB (*slr1513*) and glycogen-branching enzyme GlgB (*sll0158*) or bicarbonate transporter BicA (*sll0834*) served as positive controls (**+**), while glucose-1-phosphate adenylyltransferase (GlgC, *slr1176*) served as a negative control (Selim et al., 2021; Selim et al., 2023; Haffner et al., 2025).

### Isothermal titration calorimetry (ITC) analysis

The binding affinities of CdgR to c-di-GMP and cyclic dimeric adenosine monophosphate (c-di-AMP) were tested using isothermal titration calorimetry (ITC), as previously described (Haffner et al., 2023; Mantovani et al., 2024). ITC measurements were performed using a MicroCal PEAQ-ITC (Malvern Panalytical, Westborough, MA, USA) at 25 °C with a reference power of 10 µcal/s (Mantovani et al., 2024). For competition assays, CdgR protein was incubated with different nucleotides (ATP, ADP, AMP, cAMP, cGMP, c-di-AMP, or c-di-GMP) in the ITC cell and titrated against 1 mM c-di- AMP or c-di-GMP. Data were analyzed using one set of binding site models with the MicroCal PEAQ-ITC analysis Software (Malvern Panalytical). The dissociation constants K_D_ and ΔH values were calculated after subtracting the dilution heat from the control ITC runs.

### Differential radial capillary action of ligand assay (DRaCALA)

The binding affinities of CdgR for c-di-GMP and c-di-AMP were determined using DRaCALA, as described previously (Samir et al., 2025a). For competition binding assays, CdgR was mixed with 2 nM radioactively labeled [^32^P]c-di-AMP (∼ 6000 mCi/μmol) in a binding buffer (10 mM Tris/HCl [pH 8.0], 100 mM NaCl, and 5 mM MgCl_2_) and incubated at room temperature for 2 min. Then, 1 mM unlabeled competing nucleotides (c-di-AMP, c-di-GMP, ATP, ADP, or cAMP) were added to the reaction mixture and incubated at room temperature for 15 min. To compare the binding affinities of CdgR for c-di-GMP and c-di-AMP, titration binding assays were performed, and the IC50 values for each ligand were calculated. In the titration binding assays, unlabeled c-di-GMP or c-di-AMP (in the range of 0.1-1 mM) were titrated against a constant protein concentration incubated with 2 nM of a radioactively labeled [^32^P]c-di-AMP (∼ 6000 mCi/μmol). After incubation, 10 μL of each mixture was dropped onto a nitrocellulose membrane (AmershamTM ProtanTM 0.2 μm NC, Catalogue No10600001, Cytiva Europe GmbH, Freiburg) and left to dry. The nitrocellulose membranes were then transferred to an X-ray film cassette, and the imaging plates (BAS-IP MS 2025, 20 × 25 cm, FUJIFILM Europe GmbH, Düsseldorf, Germany) were placed directly onto the nitrocellulose membrane. The cassettes were then closed and incubated overnight. The next day, the plates were imaged using a TyphoonTM FLA9500 PhosphorImager (GE Healthcare). For [^32^P]c-di-AMP signal quantification, the image analysis software Image Studio Lite Ver 5.2.5 was used to calculate the fraction of bound nucleotides according to Samir et al. (2025a).

### In vitro pulldown analysis

The interaction/binding between CdgR and SyCRP1/SyCRP2 was tested in the presence and absence of c-di-AMP and c-di-GMP. Crude cell extracts of *E. coli* BL21 expressing CdgR-His, SyCRP1-GST, SyCRP2-GST, and SyCRP2-His were used for pulldown experiments (Supplementary Figure S1). Three mg protein of CdgR crude cell extracts was first incubated with buffer or 0.5 mM c-di-AMP, c-di-GMP for 30 min, followed by incubation with 3 mg of cell extracts expressing either SyCRP1 or SyCRP2 for 1 h to allow interaction. For the positive control, 3 mg of protein from each of the SyCRP1 and SyCRP2 crude extracts were mixed and incubated for 1 h. For each sample, 150 µL of Ni^2+^-NTA magnetic beads (Genaxxon) were washed twice with washing buffer (50 mM Tris/HCl, 200 mM NaCl; pH 7.5) before adding the crude protein mixture. The protein mixtures were then incubated with the magnetic beads for 30 min, followed by washing with washing buffer 3-5 times to remove unbound proteins. Each sample was then eluted with 20 µL of elution buffer (50 mM Tris/HCl, 200 mM NaCl, 500 mM imidazole; pH 7.5). To analyze the interaction between CdgR and SyCRP1/SyCRP2, 20 µL of the elution fraction of each sample was applied to a 12% SDS-PAGE gel, followed by western blotting. Anti-His-tag antibody at a dilution of 1:2000 was used to detect the immobilized His-tagged CdgR and SyCRP2, while anti-GST-tag antibody (10000-0-AP, Proteintech) at a dilution of 1:2000 was used to detect the GST-tagged SyCRP1-GST and SyCRP2-GST co-eluted with the immobilized CdgR. The co-elution of GST-tagged SyCRP1 with immobilized His-tagged SyCRP2 was used as a positive control.

### Protein expression and purification, and analytical size exclusion chromatography coupled to multiangle light scattering analysis (SEC-MALS)

His-tagged CdgR was expressed and purified as previously described (Samir et al., 2025a). Analytical SEC-MALS was conducted as described previously (Selim et al., 2019; Walter et al., 2019; Selim et al., 2020) to calculate the molar mass of CdgR protein and to ensure that CdgR was in its native dimeric state (Samir et al., 2025a; Samir et al., 2025b).

### Mass photometry

Mass photometry measurements were performed as described by Samir et al. (2025a). Microscope coverslips (No. 1.5H, 24 × 5, Marienfeld, Germany) were cleaned with Milli-Q water, ethanol, and isopropanol (HPLC grade) and left to dry. Silicon gaskets (3 mm diameter × 1 mm Depth, Grace Bio-Labs, Bend, OR, USA) with nine wells were rinsed with water and placed on a clean coverslip. Immersion oil was applied to the microscope objective, and the coverslip was fixed on the microscope stage. To calibrate the machine, 19 μL of buffer (50 mM Tris-HCl pH 8, 300 mM NaCl) was added to a gasket well, and the optimal focus position was identified and maintained with the 7 autofocus system. Then, 1 μL of a bovine serum albumin (BSA) / Thyroglobulin (TG) mixture at 40 nM was added to the buffer drop and mixed well with a pipette tip. The calibration video frames were recorded with an acquisition time of 60 s. Acquisition was performed following the acquisition software manual using the AcquireMP software with a TwoMP mass photometer (Refeyn, Oxford, UK). BSA can exist as a monomer (66 kDa) and/or dimer (132 kDa), whereas TG can exist as a monomer (330 kDa) and/or dimer (660 kDa). For the calibrations, at least three masses were considered to construct a standard curve. For the CdgR-SyCRP1/2 complex experiments, samples were diluted with buffer (50 mM Tris-HCl pH 8, 300 mM NaCl) to different working concentrations in the nanomolar range. The protein mixture, at a concentration of 10 nM each (considering 20 μL of final volume in the gasket drop), was incubated with c-di-AMP or c-di-GMP at the indicated concentration in each experiment and maintained at room temperature for 3 min. The mixture was then transferred to a buffer drop on the microscope coverslips previously used to find the focus. Initially, 18 μL of buffer was used to find the focus, but this volume was slightly reduced following an increase in the c-di-MNP concentration. In the absence of c-di-MNP, 18 μL of buffer was used to find the focus, and 2 μL of the protein mixture was used in the measurements. To examine the effect of cAMP on the CdgR-SyCRP1/2 complex, the proteins were incubated with cAMP before the addition of c-di-NMP. Each protein sample was measured at least twice independently, and a calibration measurement was performed before each experiment. Video files were processed using the DiscoverMP analysis software to analyze the mass events and generate graphs. All proteins, effectors, and buffers were centrifuged and filtered before the measurements.

## Results

### *Synechocystis* CdgR binds specifically c-di-GMP and with high affinity

CdgR/ComFB signaling proteins are highly conserved in bacteria and act as c-di-NMP-sensing modules that may bind to both c-di-AMP and c-di-GMP (Samir et al., 2025a). To gain insight into the binding properties of *Synechocystis* CdgR, we heterologously expressed and purified *Synechocystis* CdgR in its native dimeric state (Supplementary Figure S2). Using radiolabeled [^32^P]c-di-AMP, we performed DRaCALA assays (Roelofs et al., 2011) to test c-di-AMP binding to CdgR in competition with various unlabeled nucleotides (at 1mM), including c-di-GMP (Figure 1A). The DRaCALA binding assay revealed strong binding of [^32^P]c-di-AMP to CdgR, comparable to that of SbtB, a known c-di-AMP receptor protein in cyanobacteria (Selim et al., 2021). In the competition assays, unlabeled c-di-AMP competed with [^32^P]c-di-AMP for binding to CdgR. Other adenosine-based nucleotides (ATP and ADP), including the second messenger cyclic adenosine monophosphate (cAMP), could not compete with [^32^P]c-di-AMP for binding to CdgR (Figure 1A). Unlabeled c-di-GMP competed more efficiently than c-di-AMP with [^32^P]c-di-AMP (Figure 1A), indicating that CdgR binds to c-di-GMP. To further explore the relationship between c-di-AMP and c-di-GMP binding to CdgR, we titrated both unlabeled c-di-GMP and c-di-AMP (0.1-1 mM) to compete with [^32^P]c-di-AMP (Figure 1B). We then calculated the IC_50_, which was 20.3±5.3 µM for c-di-GMP and 126.0±21.1 µM for c-di-AMP (Figure 1B). This result implies that *Synechocystis* CdgR preferentially binds c-di-GMP.

**Figure 1:**
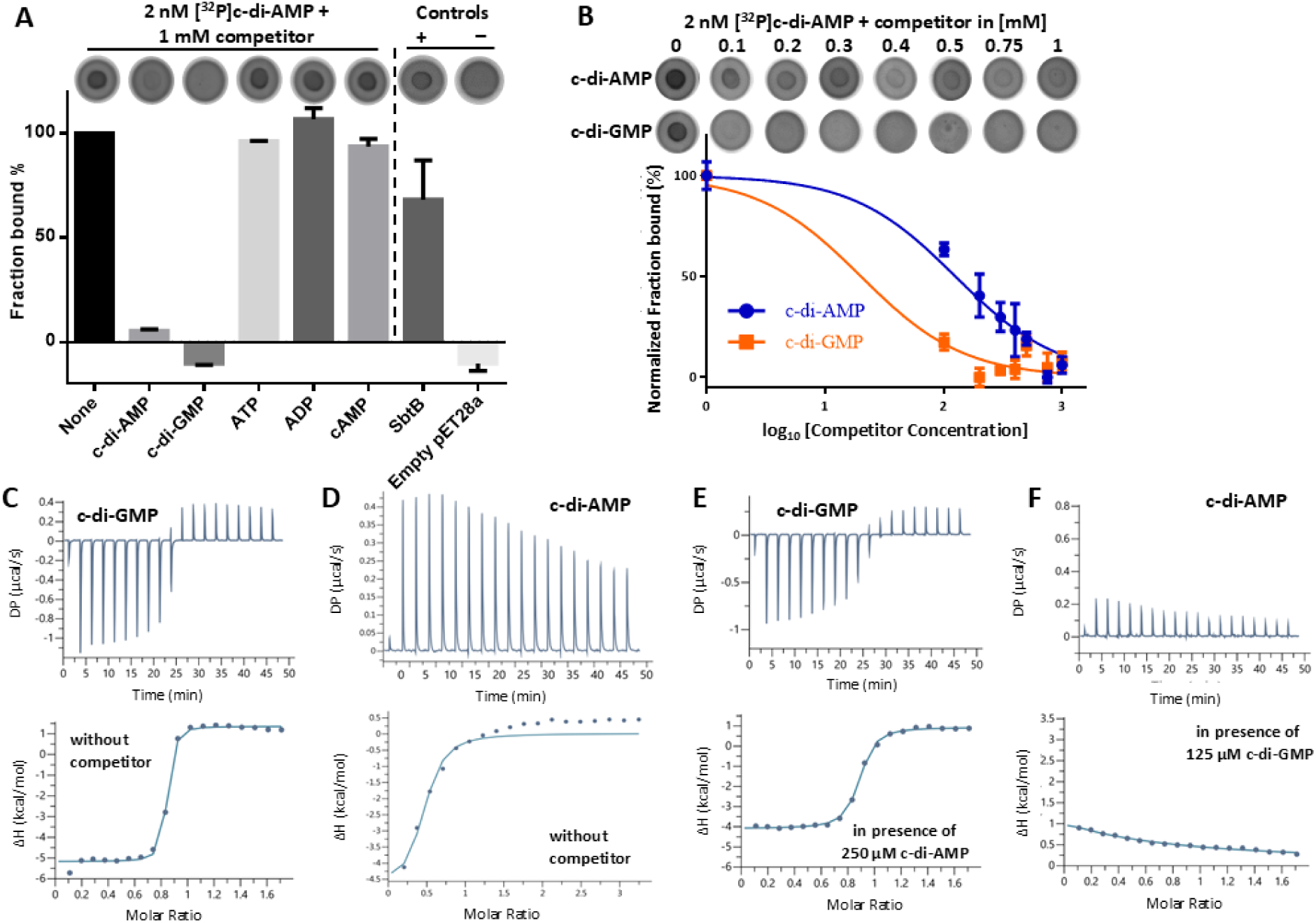
c-di-NMP binding properties of CdgR. **(A)** DRaCALA competition binding assay with CdgR showing the competitive binding of radiolabeled [^32^P]c-di-AMP with different nucleotides (1 mM) compared to the absence of competitor (none). SbtB was used as a positive control for c-di-AMP binding assays **(+)**. The fraction of bound-[^32^P]c-di-AMP to CdgR represents the mean ± SD relative to the no competitor control. **(B)** DRaCALA titration binding assay for unlabeled c-di-AMP and c-di-GMP, showing the efficiency of c-di-GMP in competing with [^32^P]c-di-AMP compared to c-di-AMP. **(C-F)** ITC analysis of c-di-GMP **(C)**, c-di-AMP **(D)**, c-di-GMP (in presence of 250 µM c-di-AMP) **(E)**, and c-di-AMP (in presence of 125 µM c-di-GMP) **(F)** binding to CdgR. Upper panels show the raw ITC data in the form of heat produced during the titration of c-di-GMP/c-di-AMP on CdgR protein; lower panels show the binding isotherms and the best-fit curves according to the one binding site model.

To confirm these results and obtain quantitative data on the binding affinities of c-di-GMP and c-di-AMP for CdgR, we performed isothermal titration calorimetry (ITC). ITC experiments revealed that c-di-GMP binds to CdgR exothermically with high affinity in the low nanomolar range (K_D_ ∼80 nM; Figure 1C). This affinity is comparable to that of other high-affinity c-di-GMP binders, such as PleD, FimX, PilZ, and MshEN (Chan et al., 2004; Navarro et al., 2009; Pultz et al., 2012; Neißner et al., 2025), including *Bacillus* ComFB (Hahn et al., 2025). In contrast, c-di-AMP binds to CdgR endothermically with a lower affinity in the micromolar range (K_D_ ∼2.9 ± 0.5 µM; Figure 1D), which further confirms that CdgR binds to c-di-GMP more specifically. At saturating concentrations of c-di-AMP (125-500 µM), c-di-GMP competes efficiently for CdgR binding, however, the binding enthalpy of c-di-GMP is slightly reduced (Table 1; compare isotherm of Figure 1E with Figure 1C and Supplementary Figure S3F), causing a 5.5-fold increase in the K_D_ value (∼440±110 nM). In contrast, the c-di-AMP binding enthalpy was strongly diminished in the presence of only 125 µM c-di-GMP (compare the isotherm of Figure 1D with Figure 1F), yielding a very high K_D_ value (∼244±32.4 µM), which is above the physiological concentration of this dinucleotide (Supplementary Figure S4) (Selim et al., 2021).

**Table 1:** Binding kinetics of c-di-GMP and c-di-AMP to CdgR.

| **Titrant** | **Competitor** | **Average $K_D$**<br>**($\mu\text{M}$)<sup>a</sup>** | **$\Delta H$ (kcal mol<sup>-1</sup>)<sup>b</sup>** | **$\Delta G$**<br>**(kcal mol<sup>-1</sup>)<sup>c</sup>** |
| --- | --- | --- | --- | --- |
| c-di-GMP | --- | $0.083 \pm 0.03$ | $-6.8 \pm 0.33$ | -9.7 |
| c-di-GMP | c-di-AMP (125-500 $\mu\text{M}$) | $0.44 \pm 0.11$ | $-4.3 \pm 0.9$ | -8.7 |
| c-di-GMP | 150 $\mu\text{M}$ ATP | $0.09 \pm 0.05$ | $-5.0 \pm 0.15$ | -9.6 |
| c-di-GMP | 150 $\mu\text{M}$ ADP | $0.5 \pm 0.15$ | $-5.4 \pm 0.15$ | -8.6 |
| c-di-GMP | 150 $\mu\text{M}$ AMP | 4.2 | -5.7 | -7.3 |
| c-di-GMP | 150 $\mu\text{M}$ cAMP | $0.3 \pm 0.1$ | $-4.8 \pm 0.14$ | -8.9 |
| c-di-GMP | 150 $\mu\text{M}$ cGMP | $0.8 \pm 0.4$ | $-5.7 \pm 0.3$ | - 8.3 |
| c-di-AMP | ---- | $2.9 \pm 0.5$ | $4.77 \pm 0.05$ | -7.3 |
| c-di-AMP | 125 $\mu\text{M}$ c-di-GMP | $244 \pm 32.4$ | 80.0 | -4.9 |

The raw ITC data were fitted using a one-site binding model for the monomeric CdgR protein.
a) The dissociation constant ( $K_D$ ) values correspond to the mean ± SD. b) Enthalpy; c) Gibbs free energy. To test potential competition by other nucleotides, the protein was preincubated with the indicated “competitor, followed by titration with c-di-GMP or c-di-AMP as indicated.

To further explore the relationship between c-di-GMP binding and the binding of other, more abundant nucleotides (ATP, ADP, and AMP), and other second messengers (cGMP and cAMP), we examined the ability of CdgR to bind c-di-GMP in the presence of saturating concentrations of these nucleotides (150 µM). The presence of ATP did not significantly affect c-di-GMP binding to CdgR, whereas ADP, cAMP, or cGMP slightly reduced the affinity of c-di-GMP for CdgR in a manner similar to that of c-di-AMP (Table 1; Supplementary Figure S3). Moreover, the binding enthalpy and Gibbs free energy (ΔG) for c-di-GMP-binding were reduced in the presence of AMP (Table 1; compare the isotherm in Figure 1C with Supplementary Figure S3D), causing an increase in the K_D_ value.

To further exclude any possible role of c-di-AMP binding to CdgR (Figure 1), we checked the relative intracellular concentrations of c-di-GMP and c-di-AMP under white light illumination. The intracellular concentration of c-di-GMP was almost 11 times higher than that of c-di-AMP (Supplementary Figure S4), suggesting that CdgR mainly binds c-di-GMP in a cellular context.

### CdgR is involved in type IV pilus dependent functions in *Synechocystis*

To study the effect of the lack of *cdgR* on the behavior of *Synechocystis* cells, we replaced the entire gene with a kanamycin resistance cassette (Supplementary Figure S5) and assessed the phototactic motility of two different clones on soft agar plates under white and blue light. Under lateral illumination with white light, the *cdgR::Km* mutant displayed enhanced motility compared to the wild-type background strain (Figure 2A), suggesting that *Synechocystis* CdgR plays an inhibitory role in phototaxis. Single-cell motility examined under the microscope using unilateral red light revealed that *cdgR::Km* mutant cells did not move at a higher speed, but rather moved more directionally towards the light source (Figure 2B-C). Previous studies have shown that under blue light, Cph2-dependent synthesis of c-di-GMP leads to the inhibition of motility (Savakis et al., 2012) and aggregation of cells (Conradi et al., 2019). However, motility was still inhibited in the *cdgR::Km* mutant under blue light compared to that in the wild-type strain (Figure 2D). In contrast to its effect on motility, deletion of *cdgR* did not alter the aggregation phenotype (Figure 2E), suggesting that CdgR is not directly involved in this process. A previous study showed that in *Synechocystis,* elevated c-di-GMP levels result in the downregulation of the *pilA5*-*pilA6* gene cluster, which encodes minor pilins (Wallner et al., 2020). Given the critical role of PilA5 in natural competence in *Synechocystis* (Oeser et al., 2021), we assessed the ability of the *cdgR::Km* mutant cells to be transformed by exogenous DNA. Deletion of *cdgR* resulted in a nearly complete loss of natural competence suggesting that DNA uptake by IV pili is impaired (Figure 2F). This result also confirmed previous studies by Samir et al. (2025a). However, the *cdgR* mutant is able to move and to aggregate, two function which depend on type IV pili. Thus, CdgR is involved in some, but not all, type IV pilus-related functions. This indicates that pilus biogenesis is largely unaffected in the *cdgR::Km* mutant. In addition, deletion of *cdgR* did not alter intracellular c-di-GMP levels (Supplementary Figure S4). Together, these findings suggest that CdgR specifically regulates a subset of type IV-pilus-dependent functions rather than affecting pilus assembly or global c-di-GMP signaling.

**Figure 2:**
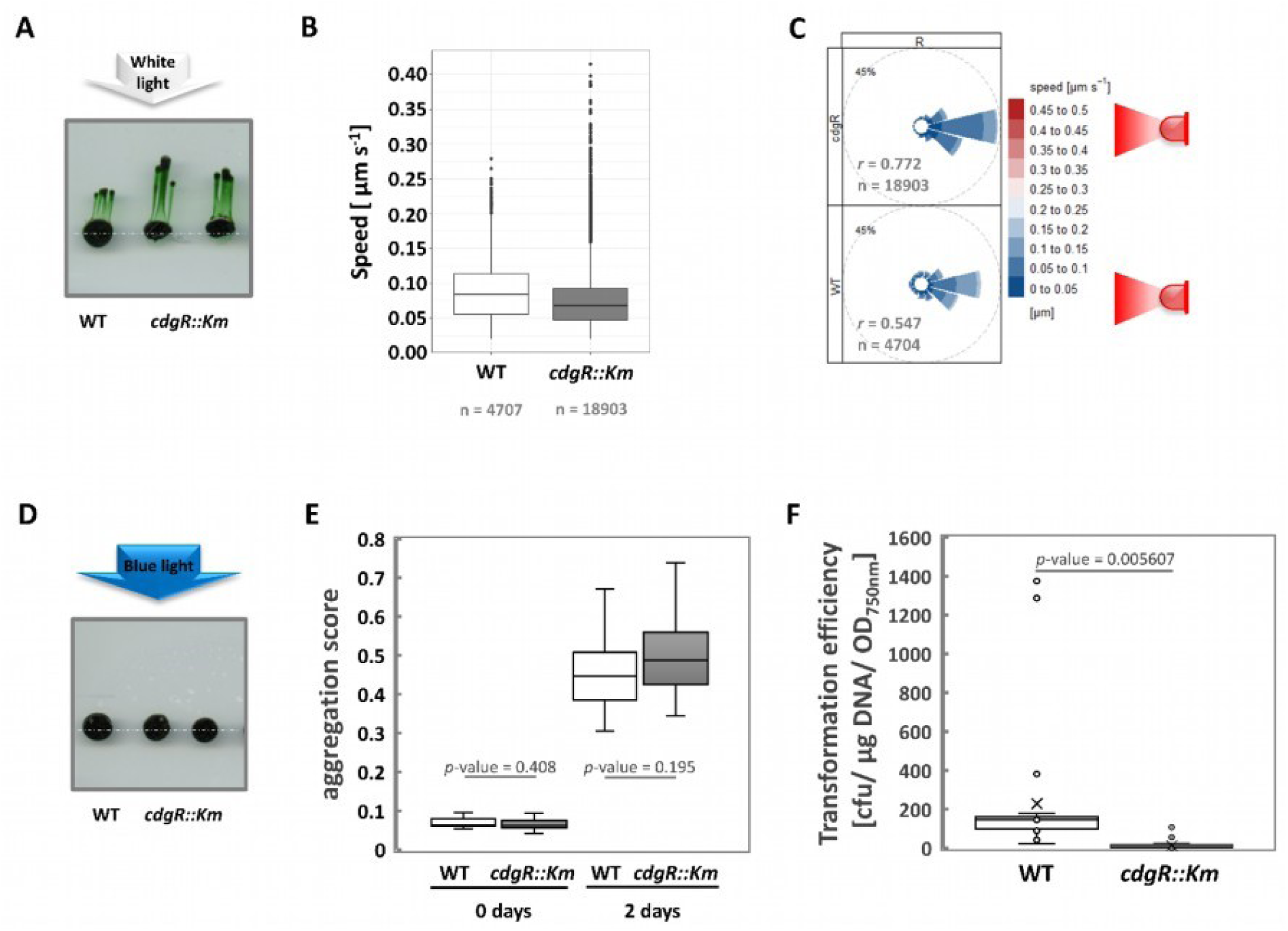
Phenotypes of the *cdgR::Km* mutant strain. **A:** Phototaxis assays of the *cdgR::Km* mutant strain in response to white light. *Synechocystis* wild-type and *cdgR::Km* mutant strains were cultured on BG11 agar plates (0.5% (w/v)) supplemented with 11 mM glucose and 10 mM TES buffer (pH 8.0). The cells were then unidirectionally illuminated with 5 μmol photons m^−2^ s^−1^ of white light. Images were captured after 14 days of incubation. The white dashed lines indicate the initial spotting areas. **B and C:** Single-cell motility of the *cdgR::Km* mutant strain. Cells of *Synechocystis* wild type and the *cdgR::Km* mutant were illuminated with unidirectional red light (40 µmol photons m^-2^ s^-1^) on BG11 agarose plates under a microscope, and the displacement of the cells over a 3 min time-frame was captured 2 min after the onset of illumination. Raw tracks of moving cells were determined using Fiji’s TrackMate plugin. The velocity and directionality of the moving cells were analyzed using R software. **B:** Mean speed of wild-type and *cdgR::Km* cells. **C:** The mean resultant length from a Rayleigh test (*r*) and the number of tracked motile cells (*n*) are shown. **D:** Phototaxis assays of the *cdgR::Km* mutant strains in response to 5 µmol photons m^-2^ s^-1^ of blue light. **E:** Flocculation assay of the *cdgR::Km* strain under white light. Aggregation values of *Synechocystis* wild type (white boxes, *n* = 12) and *cdgR::Km* (grey boxes, *n* = 24) are displayed before and after two days of incubation. A two-sample *t*-test assuming equal variances was applied **F:** Transformation efficiency of the *cdgR::Km* mutant strain. These experiments determined the transformation efficiency of *Synechocystis* wild-type (n = 24) and the *cdgR::Km* mutant strains (n = 32). The number of colony-forming units (cfu) was counted after 12 days, and the transformation efficiency was normalized to the optical density at 750 nm (OD_750nm_) and the amount of DNA used. For each transformation, one microgram of pJET*-ΔcrhR*-SpecR plasmid DNA was used, conferring spectinomycin resistance. A Welch’s *t*-test was applied. Wild type, WT.

The *cdgR* gene forms a transcriptional unit with *hfq* (Figure S5A) (Kopf et al., 2014), which is involved in type IV pilus functions in *Synechocystis* (Dienst et al., 2008). *Hfq* is the first open reading frame in this operon and an Δ*hfq* mutant is non-transformable (Dienst et al., 2008). This synteny between *hfq* and *cdgR* is conserved in several unicellular cyanobacteria (Samir et al., 2025a). To exclude an effect of *cdgR* mutation on Hfq function, we complemented the phenotype by expressing a FLAG-tagged *cdgR* gene from its own promotor on a self-replicating plasmid (Supplementary Figure S6). Transformability was completely restored in this strain (Supplementary Figure S6D). To further assess putative polar effects of the integration of the kanamycin cassette in this region, we generated a new *cdgR* knockout strain (Δ*cdgR*) using the CRISPR-Cpf1 based technique (Niu et al., 2019). The resulting mutant had an in-frame deletion of *cdgR* (Supplementary Figure S7) and showed enhanced movement similar to that of *cdgR::Km,* compared to the wild-type (Figure 3A). Ectopic expression of *cdgR* from a neutral locus (C-Δ*cdgR*) restored slower wild-type movement (Figure 3A). These results suggest that the enhanced phototaxis of both mutant strains mainly depended on the lack of *cdgR*.

**Figure 3.**
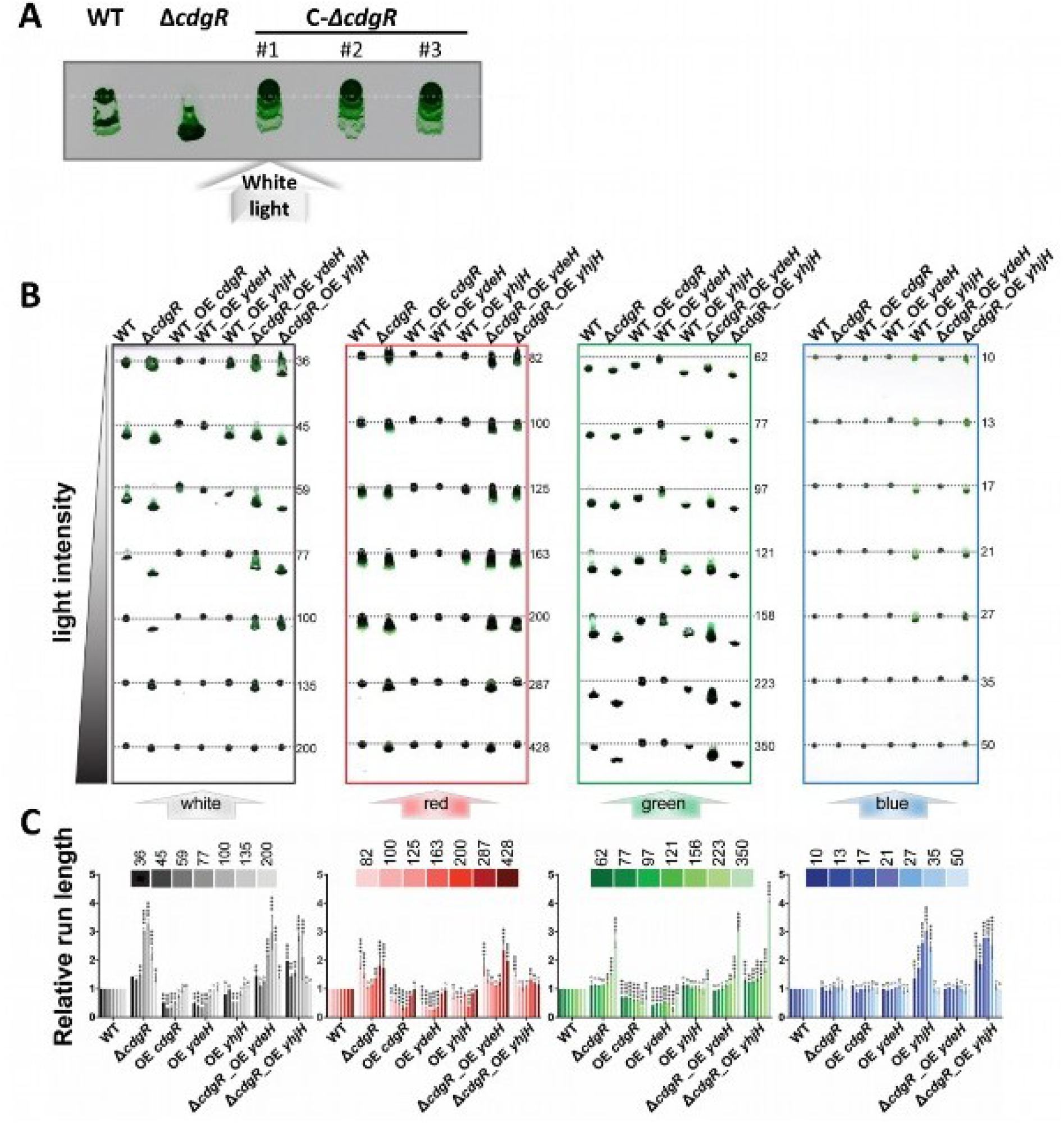
Regulation of phototaxis by CdgR in *Synechocystis* depends on light intensity and quality. **A:** Phototaxis assays of the Δ*cdgR*(Cfp1) mutant strain in response to white light. *Synechocystis* wild type, Δ*cdgR*(Cfp1) and complementation strain (C-Δ*cdgR*) were cultivated on BG11 agar plates (0.7% (w/v)) supplemented with 10 mM glucose and 12.5 mM TES buffer (pH 8.0). The cells were then unidirectionally illuminated with 5 μmol photons m^-2^ s^-1^ of white light for four days. The dashed lines indicate the initial spotting areas. Three independently generated complementation clones (#1-3) were used. **B:** BG11 agar (0.7% (w/v)) supplemented with 10mM glucose and 12.5 mM TES buffer (pH 8.0) was filled in a (24 cm x 24 cm) square plates for phototaxis assay. All the indicated strains were inoculated on the plate at different distances from the white, red, green or blue LED light source, which was placed laterally to provide unidirectional illumination. The position of the initial spot was marked by grey dashed lines and the light intensity (μmol photons m^-2^ s^-1^) was specified by the number on the right. **C:** Quantification of the relative run length from phototaxis assay from (**B);** light intensity is as indicated. Data are shown as mean ± SD (n =2). Significant differences compared to the wild type were analyzed using two-way ANOVA. Significance thresholds: * p < 0.05, ** p < 0.01, *** p < 0.001, **** p < 0.0001; n, not significant.

To explore whether the observed enhanced movement of the mutants depends on light quality, we compared the phototactic behavior of the Δ*cdgR* mutant and the wild type under different light qualities and intensities (blue, green, and red). Cells spotted at different positions on a large square agar plate along the light direction experienced different light intensities (Figure 3B). Interestingly, we observed that the accelerated motility phenotype of the Δ*cdgR* mutant was more pronounced not only under a specific intensity range of white light but also under red and particularly green light (Figure 3B). Quantification of the phototactic motility phenotype by normalizing the distance moved by each mutant to that of the wild type under the same conditions revealed distinct patterns under different light conditions. Under white light, a notable increase in the distance moved occurred for the Δ*cdgR* mutant between 77-100 µmol photons m^-2^ s^-1^. Below this range, Δ*cdgR* cells exhibited only a slight increase in distance compared to wild-type cells, whereas above this range, the movement of both strains was inhibited by high light intensity in a comparable manner (Figure 3C). Under red light, enhanced motility of Δ*cdgR* was observed at low (<100 µmol photons m^-2^ s^-1^) and high (>163 µmol photons m^-2^ s^-1^) light intensities (Figure 3C). Green light enhanced cell motility in Δ*cdgR,* especially at intensities above 121 µmol photons m^-2^ s^-1^, even at the highest intensity of 350 µmol photons m^-2^ s^-1^, which completely inhibited cell movement of the wild type (Figure 3C). In contrast, the overexpression of *Synechocystis* CdgR (WT_OE *cdgR* strain) resulted in significantly reduced motility across a broad range of light intensities under white, red, and green light when compared to the wild type (Figure 3B and 3C), indicating that CdgR is involved in the inhibition of cell motility under these conditions.

To determine whether the *cdgR* mutation caused any defects in pilus formation, we used electron microscopy to observe the cell surface structures. Our results showed that both thick and thin pili were present in the Δ*cdgR* mutant, and no obvious differences in cell surface structures were observed between the mutant and wild-type strains (Supplementary Figure S8).

### C-di-GMP dependent phototaxis control

As CdgR is the first c-di-GMP receptor protein found in *Synechocystis*, we manipulated cellular c-di-GMP levels by overexpressing *E. coli ydeH* or *yhjH* genes in *Synechocystis* to further investigate the role of c-di-GMP in CdgR-mediated motility regulation. The *ydeH* and *yhjH* genes encode a c-di-GMP synthetase and a c-di-GMP hydrolase, respectively (Boehm et al., 2009). These two genes are frequently used to manipulate c-di-GMP levels in different heterologous hosts (Savakis et al., 2012; Enomoto et al., 2015; Huang et al., 2021). The two genes were expressed in both the wild type (WT_OE *ydeH* and WT_OE *yhjH*) and the Δ*cdgR* mutant (Δ*cdgR*_OE *ydeH* and Δ*cdgR*_OE *yhjH*), respectively, using a chimeric promoter regulated by copper and theophylline. As expected, WT_OE *ydeH*, which has a predicted high c-di-GMP level in the wild-type background, showed inhibition of phototactic motility under white, red, and green light (Figure 3B). This is similar to the effect observed in the *cdgR* overexpressor strain (WT_OE *cdgR*), as quantified by the colony migration distance (Figure 3C). These results are consistent with the inhibitory role of c-di-GMP in cell motility (Savakis et al., 2012). Conversely, the Δ*cdgR*_OE *ydeH* mutant, which had a high level of c-di-GMP in the absence of CdgR, was unable to inhibit motility compared with the WT_OE_*ydeH* mutant (Figure 3B). It exhibited a migration distance similar to that of the Δ*cdgR* mutant (Figure 3C). Therefore, the inhibitory effect of c-di-GMP on phototaxis is mediated by CdgR. The WT_OE *yhjH* overexpressing strain exhibited an inhibitory effect similar to that of WT_OE *cdgR* and WT_OE *ydeH* under white and red light but exhibited a wild-type-like phenotype under green light (Figure 3B). However, the strain that overexpressed *yhjH* in the absence of CdgR (Δ*cdgR*_OE *yhjH*) exhibited a Δ*cdgR*-like phenotype (Figure 3B). Taken together, these findings suggest that c-di-GMP-mediated motility inhibition requires CdgR under white, red, and green light conditions. Blue light stimulates c-di-GMP synthesis due to the light-dependent diguanylate cyclase activity of the photoreceptor Cph2. This blue-light dependent c-di-GMP increase inhibits phototaxis in wild-type cells (Fiedler et al., 2005; Savakis et al., 2012; Wallner et al., 2020). Due to a lack of this c-di-GMP increase in the *cph2* mutant, the cells therefore, move towards blue light, in contrast to the wild type. If CdgR acts as the sole sensor for the c-di-GMP produced under blue-light conditions, a *cdgR* mutant should exhibit a phototactic phenotype similar to that of the *cph2* mutant. However, this was not what we observed. Notably, under blue light, none of the mutants moved, except for two mutants that expressed the c-di-GMP-degrading enzyme YhjH. Furthermore, a Δ*cdgR*Δ*cph2* double mutant displayed a *cdgR::Km*-like hypermotile phenotype under white light, but a Δ*cph2*-like phenotype under blue light (Supplementary Figure S9). In contrast to the strict dependency on CdgR under red and green light, the c-di-GMP-mediated inhibition mechanism under blue light operates independently of CdgR, implying that CdgR is the dominant regulator under white light while Cph2 plays a predominant role under blue light.

### CdgR regulates the expression of minor pilin genes

A previous study showed that an elevation in c-di-GMP levels resulted in changes in motility-related genes (Huang et al., 2021). Furthermore, *Anabaena* CdgR interacts with the global transcription factor DevH (Zeng et al., 2023). Therefore, we investigated the impact of the *cdgR* mutation on the transcriptome using the Δ*cdgR* mutant strain in comparison to the wild-type. Cells were grown on phototaxis plates under conditions where the Δ*cdgR* strain exhibited enhanced motility. Transcriptomic analysis revealed only 12 differentially expressed genes (DEGs) in the Δ*cdgR* strain, comprising five upregulated and seven downregulated genes (Figure 4A, Table 2, and Supplementary Table 4). Notably, the most significantly upregulated genes, *slr1971* and *hfq (ssr3341*), are located adjacent to *cdgR,* with *hfq* in the same transcriptional unit. The gene *slr1971,* which encodes a putative peptidase, is transcribed from its own promoter downstream of *cdgR* (Kopf et al., 2014). Although we used the marker-less Δ*cdgR* strain for transcriptome analysis, these changes can still be due to the polar effects of the mutation, for example, by stabilizing the shorter *hfq* transcript.

**Figure 4:**
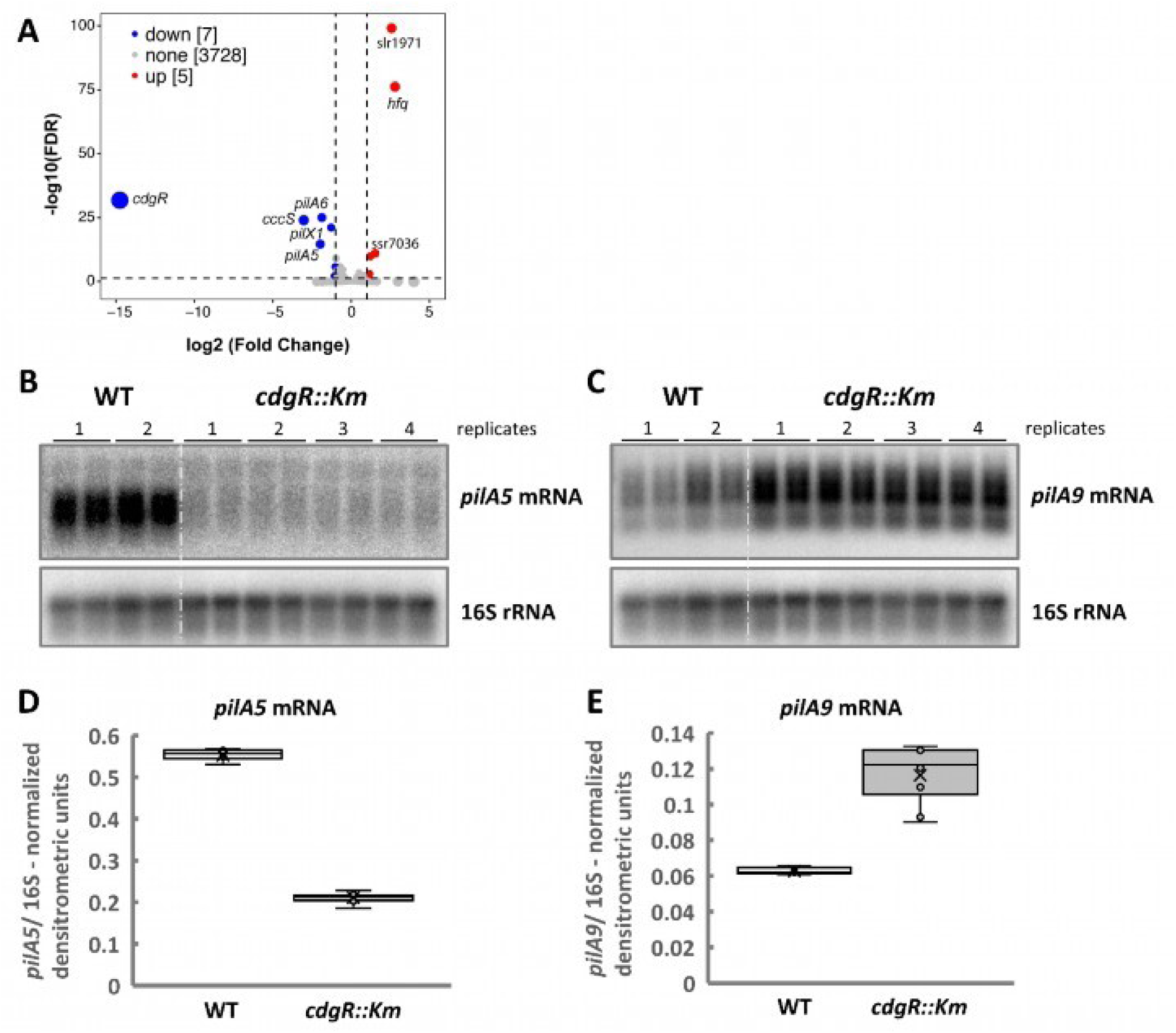
Transcriptome analysis of the *cdgR* mutant strain. **A:** Volcano plot showing the differentially expressed genes (DEGs) in the *ΔcdgR* mutant compared to the wild type determined by RNA-seq. DEGs were defined as genes with [log_2_ fold change] >1 and log_10_(FDR)<0.05. **B-E:** Expression of minor pilin genes in the *cdgR::Km* mutant strain. The *cdgR::Km* mutant strain was examined for the expression of minor pilin genes. After 24 h of exposure to white-light illumination (75 µmol photons m^-2^ s^-1^), total RNA was extracted from cells grown in BG11 medium. Three micrograms of RNA were hybridized with radioactively labeled RNA probes targeting the *pilA5* mRNA (**B**) and the 5’-UTR of the *pilA9* mRNA (**C**). A double-stranded DNA probe that hybridized with *Synechocystis* 16S rRNA was used as a loading control. Normalized mRNA expression levels of *pilA5* (**D**) and *pilA9* (**E**) were determined by densitometric quantification relative to 16S rRNA levels. Two biological replicates, each with two technical replicates, were performed for the WT. Four biological replicates, each with two technical replicates, were used for the *cdgR::Km* mutant experiment.

**Table 2:**
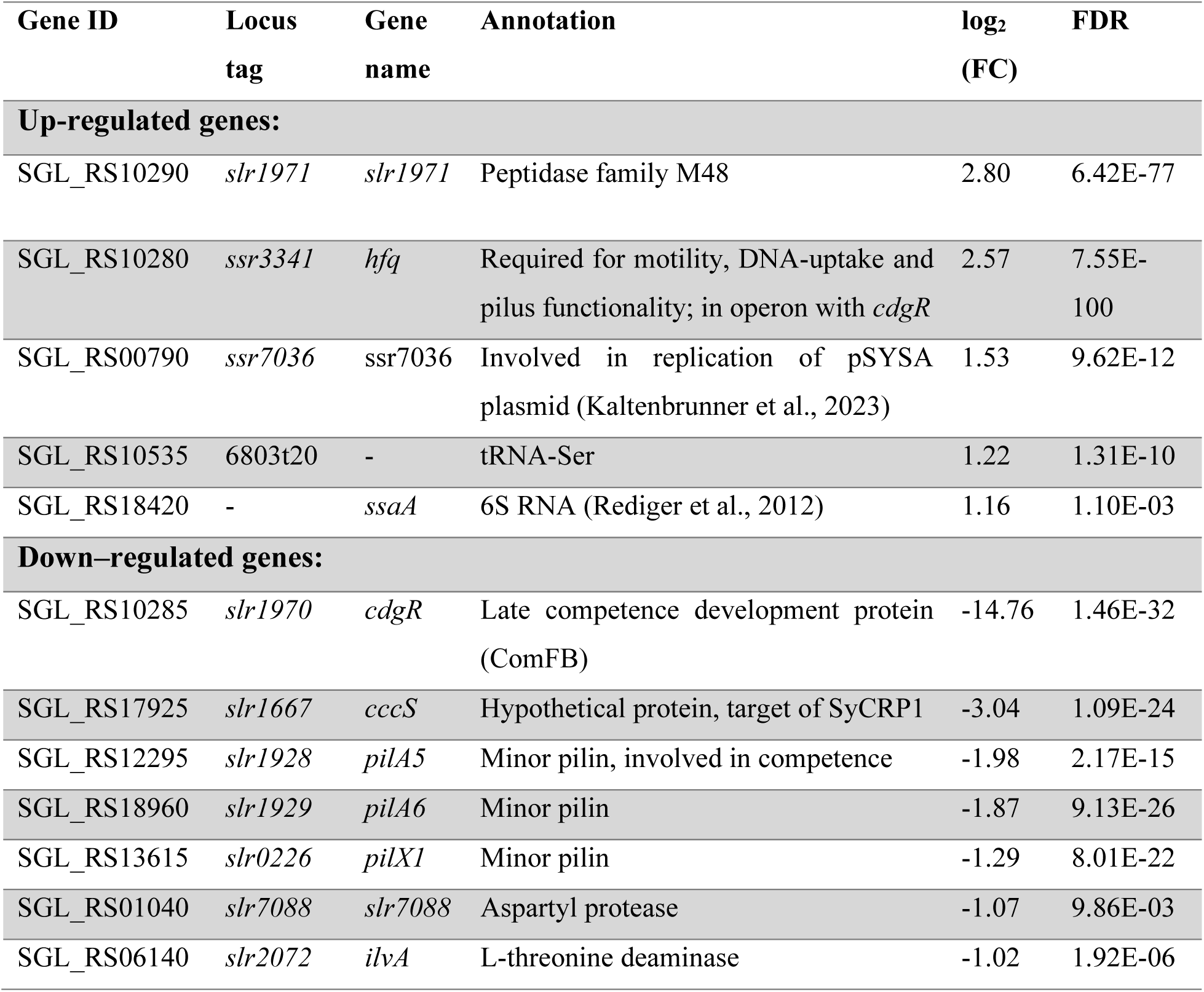
List of DEGs identified in the transcriptome analysis of the Δ*cdgR* strain. Criteria: genes with a log_2_ fold change>1 or <1 and FDR < 0.05.

Interestingly, RNA-seq analysis revealed a decrease in the abundance of the *pilA5-pilA6* mRNA, which could explain the observed defect in the natural transformation of *cdgR::Km* mutant (Figure 2F). To verify these results and to demonstrate that the minor pilin genes were also differentially regulated in the *cdgR::Km* strain, we extracted total RNA from both the wild type and *cdgR::Km* mutant strain grown in liquid culture for northern blot analysis, using a *pilA5* mRNA. Northern blot analysis revealed that, in accordance with the RNA-seq data, the transcription level of the *pilA5* gene in the *cdgR* mutant was downregulated (Figure 4B), accounting for only approximately 38% of the transcript level in the wild-type strain (Figure 4D). This suggests that CdgR regulates natural competence by modulating *pilA5-pilA6* gene expression in *Synechocystis.* Notably, when the transcriptional regulation of *pilA5-pilA6* was observed, the transcript accumulation of another minor pilin operon, *pilA9-pilA12*, was frequently altered in the opposite way (Wallner et al., 2020; Oeser et al., 2021). These minor pilins are involved in cell aggregation and motility (Oeser et al., 2021). Northern blot analysis revealed a substantial accumulation of *pilA9* (Figure 4C), with an approximately 1.85-fold increase in transcript levels compared to those in the wild-type strain (Figure 4E). RNA-seq also revealed a slight increase in *pilA9-pilA12* transcript accumulation, although the expression changes were less pronounced and not significant in this analysis (Table 2 additional information). Increased *pilA9-pilA12* expression may contribute to the accelerated motility of *cdgR* mutant cells. However, this did not extend to cell-cell aggregation, which also depends on the minor pilins PilA9-PilA12, as we observed comparable aggregation scores between the wild-type and mutant strains (Figure 2E).

### *Synechocystis* CdgR interacts with the transcription factors SyCRP1 and SyCRP2

In addition to the upregulation of the *pilA9-pilA12* operon, our transcriptome analysis (Table 2) also revealed the downregulation of *cccS*, which encodes a subunit of the putative chaperone-usher system. The *pilA9-pilA12* operon and *cccS* are known targets of the cAMP-dependent transcription factor SyCRP1 (Yoshimura et al., 2002). *Synechocystis* also encodes a second SyCRP1 homologue devoid of cAMP-binding capacity, known as SyCRP2. SyCRP2 has been implicated in the regulation of pilus biogenesis, particularly the *pilA9-pilA12* operon, through its interaction with SyCRP1 (Song et al., 2018). Additionally, we previously demonstrated that the transcription factor DevH of the CRP-family controls cell size in the multicellular cyanobacterium *Anabaena* through its interaction with *Anabaena* CdgR (Zeng et al., 2023). These findings suggest that SyCRP1 and/or SyCRP2 are likely candidates to interact with *Synechocystis* CdgR. To examine the possible interactions between CdgR and SyCRP1 or SyCRP2, we used our established bacterial adenylate cyclase two-hybrid (BACTH) interaction assay (Karimova et al., 1998; Battesti and Bouveret, 2012). In this assay, the T25 subunit of *Bordetella pertussis* adenylate cyclase (Cya) was fused to either the N- or C-terminus of CdgR, and the T18 subunit of Cya was fused to either the N- or C-terminus of SyCRP1 or SyCRP2 (Supplementary Figure S10A-B). Known SbtB interactions with GlgB (glycogen branching enzyme) or BicA (Na^+^-dependent HCO_3_^-^ transporter) were used as positive controls (Selim et al., 2021; Haffner et al., 2025; Samir et al., 2025a), and GlgC (glucose-1-phosphate adenylyltransferase) was used as a negative control (Selim et al., 2023). We also checked for the known SyCRP1-SyCRP2 interaction by fusing the T25 subunit C-terminally to SyCRP1 or SyCRP2 (Song et al., 2018). Clear interactions were observed between CdgR-SyCRP1 and CdgR-SyCRP2 for all tested combinations on both X-Gal and MacConkey agar plates (Supplementary Figure S10A-B). We also observed a clear interaction between SyCRP1-SyCRP2, which validated our BACTH approach. These results strongly support the specificity of CdgR interaction with SyCRP1 and SyCRP2.

To validate the specificity of the interaction between CdgR and SyCRP1 or SyCRP2 and to examine the influence of c-di-NMP on these interactions using an independent method, we examined the physical interaction between CdgR and SyCRP1 or SyCRP2 using *in vitro* pulldown assays (Supplementary Figure S10C-D). In the absence of c-di-GMP or c-di-AMP, CdgR interacted with SyCRP1 and SyCRP2, as indicated by the co-elution of SyCRP1 or SyCRP2 with CdgR (Supplementary Figure S10C-D). Addition of c-di-GMP seemed to weaken the CdgR-SyCRP1 and CdgR-SyCRP2 complex formation. These results are consistent with the BACTH assay data and further support our conclusion regarding the specificity of the CdgR-SyCRP1 and CdgR-SyCRP2 interactions.

### Influence of different effector molecules on CdgR-SyCRP1/2 complex formation

To gain quantitative insight into the sensitivity of the CdgR-SyCRP1 and CdgR-SyCRP2 complexes to c-di-GMP compared to c-di-AMP and to test the influence of cAMP (a known SyCRP1 effector molecule) on the assembly of the complexes, we used mass photometry. Mass photometry utilizes physiological nanomolar concentrations of proteins, therefore, it could reflect the *in vivo* situations. In the absence of c-di-GMP, CdgR formed stable complexes with SyCRP1 (Figure 5A) or SyCRP2 (Figure 6A), which dissociated efficiently in the presence of 1 or 0.5 mM c-di-GMP (Figure 5B and 6B; and Supplementary Figure S11B). Unlike c-di-GMP, c-di-AMP did not strongly affect the CdgR-SyCRP1 complex, where the complex partially dissociated at 1 mM (Figure 5B). A c-di-GMP titration experiment revealed that the CdgR-SyCRP1 complex was stable at low c-di-GMP concentrations (Figure 5C) and required at least 0.5 mM c-di-GMP to be completely disrupted (Supplementary Figure S11B). For the CdgR-SyCRP2 complex, 50-250 µM c-di-GMP was sufficient to dissociate the complex (Figure 6C and Supplementary Figure S11D), while at least 0.5 mM c-di-AMP was required to start the dissociation of CdgR-SyCRP2 complex (Figure 6B and Supplementary Figure S11C). These results further confirm the specificity of c-di-GMP binding to CdgR (Table 1; Figure 1). In contrast, c-di-AMP appears to be inefficient, requiring higher concentrations to disturb the assembly of CdgR-SyCRP1/2 complexes (Supplementary Figure S11A and S11C).

**Figure 5:**
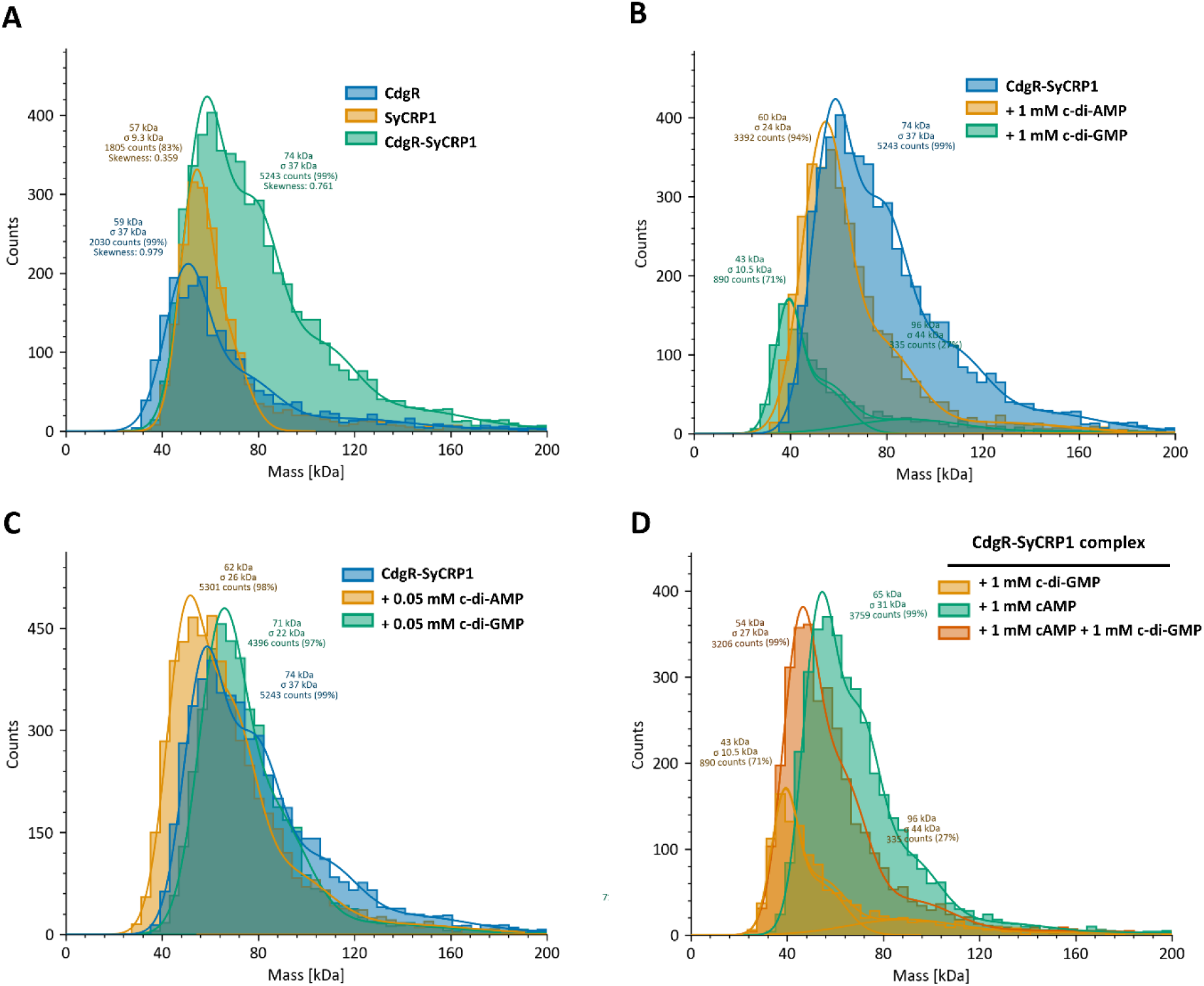
Mass photometry analysis showing the interaction between CdgR and SyCRP1. **(A).** The effect of high concentration **(B)** and low concentration **(C)** of c-di-GMP and c-di-AMP on the CdgR-SyCRP1 complex; and the effect of cAMP on the CdgR-SyCRP1 complex in absence or presence of c-di-GMP **(D)**. The complex is indicated by higher mass of 90-120 kDa.

**Figure 6:**
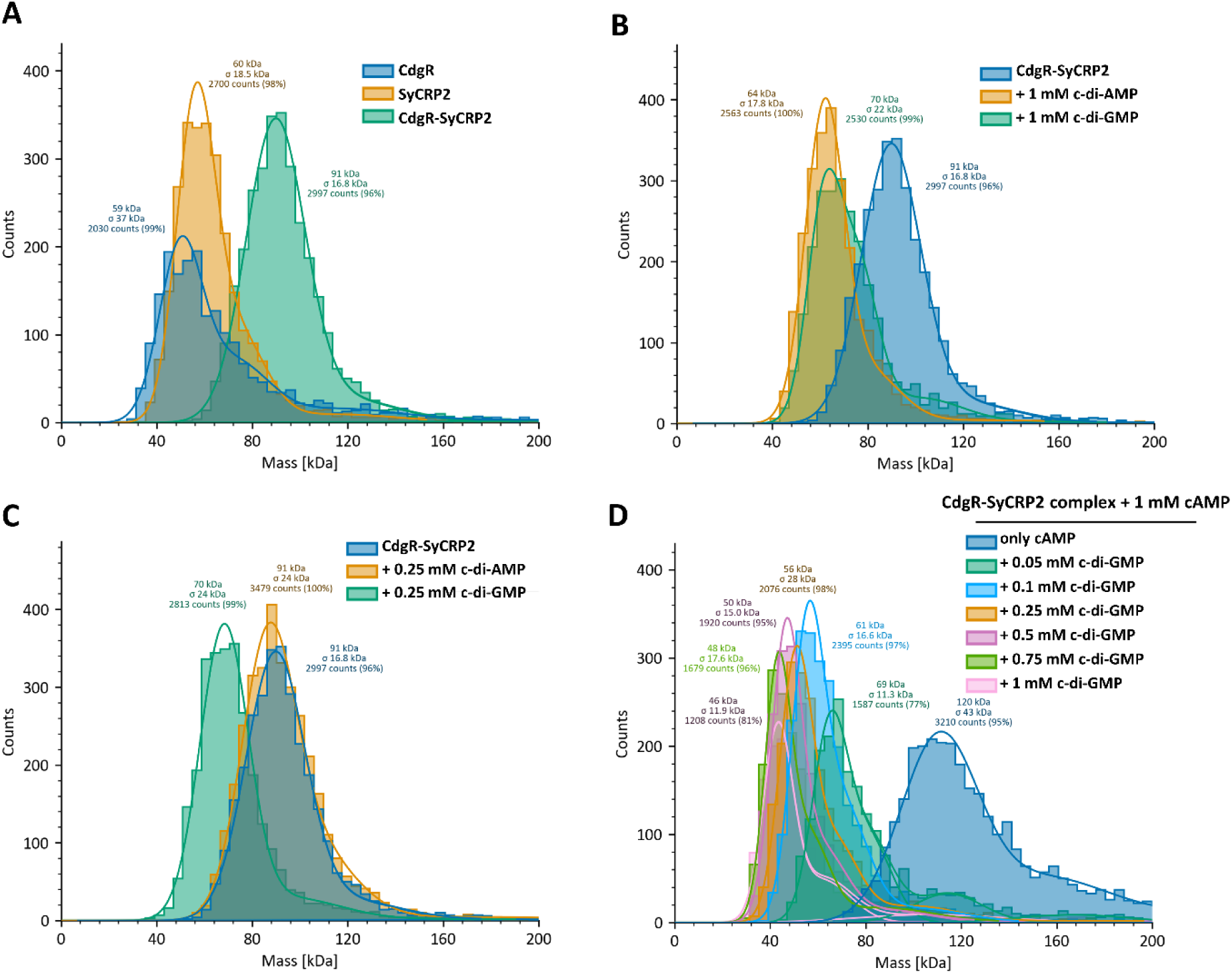
Mass photometry analysis showing the interaction between CdgR and SyCRP2. **(A).** The effect of high concentration **(B)** and low concentration **(C)** of c-di-GMP and c-di-AMP on the CdgR-SyCRP2 complex; and the effect of cAMP on the CdgR-SyCRP2 complex in absence and presence of different concentration of c-di-GMP **(D)**. The complex is indicated by higher mass of 90-120 kDa.

Unlike SyCRP1, which is a cAMP-dependent transcription factor, SyCRP2 does not bind cAMP. However, they work together to regulate the pilus machinery transcriptionally (Song et al., 2018). To examine the possibility of a crosstalk between cAMP and c-di-NMP signaling in the CdgR-SyCRP1/2 interaction, we studied the influence of cAMP on complex assembly and examined the ability of c-di-GMP or c-di-AMP to disrupt the complexes in the presence or absence of cAMP. The presence of cAMP (1 mM) did not affect the CdgR-SyCRP1/2 interaction, and the complexes formed as they did in the absence of cAMP (Figure 5D and 6D). Surprisingly, cAMP had a protective effect against c-di-GMP for the CdgR-SyCRP1 complex, preventing dissociation even at 0.5 or 1 mM of c-di-GMP (Figure 5D and Supplementary Figure S11E). In contrast, cAMP did not have a protective effect on the CdgR-SyCRP2 complex, which responded normally to c-di-NMP (Figure 6D and Supplementary Figure S11F). This is consistent with the inability of SyCRP2 to bind cAMP.

## Discussion

Previous studies have shown that the expression of several minor pilins is regulated by the levels of cAMP and c-di-GMP (Yoshimura et al., 2002; Wallner et al., 2020). cAMP is a key second messenger that regulates multiple cellular processes in both prokaryotes and eukaryotes (Gancedo, 2013). In cyanobacteria, cAMP signaling is essential for coordinating and fine-tuning cell motility, particularly in response to environmental cues (Ohmori et al., 1992; Terauchi and Ohmori, 1999,2004). In *Synechocystis*, two cAMP receptor proteins, SyCRP1 and SyCRP2, have been identified as key transcription factors involved in cell motility. While SyCRP1 directly binds to cAMP, SyCRP2 lacks conserved cAMP-binding sites (Yoshimura et al., 2000). Intriguingly, disruption of either SyCRP1 or SyCRP2 results in motility defects due to reduced pilus formation (Yoshimura et al., 2002; Bhaya et al., 2006; Song et al., 2018). It has been proposed that SyCRP2 forms a heterodimer with cAMP-bound SyCRP1 to initiate the transcription of the minor pilin gene cluster *pilA9*- *pilA12*, which is essential for cell motility (Song et al., 2018; Wallner et al., 2020).

In our model (Fig. 7), CdgR interacts with the two transcription factors SyCRP1 and SyCRP2 under low c-di-GMP conditions. At low c-di-GMP concentrations, CdgR binds to both SyCRP1 and SyCRP2, resulting in the accumulation of *pilA5-6* mRNA and basal expression of *pilA9-12*. In contrast, high c-di-GMP concentrations disrupt CdgR-SyCRP1/2 complexes. Consequently, free SyCRP2 represses the transcription of *pilA5-pilA6* and interacts with SyCRP1 to activate *pilA9-12* expression. The lack of CdgR mirrors high c-di-GMP conditions with respect to the expression patterns of the two minor pilin operons (*pilA5-6* and *pilA9-12*) as both SyCRP1/2 are in the unbound state. This observation is consistent with the phenotypes of the Δ*cdgR* strain. When the *pilA5-pilA6* operon is not expressed, the cells are not transformable, similar to the Δ*pilA5-pilA6* strain (Oeser et al., 2021). Whereas, the stronger positive phototaxis of *cdgR* mutants can be attributed to elevated expression of *pilA9-pilA12*. However, high c-di-GMP concentrations, as observed under blue light incubation, inhibit motility (Savakis et al., 2012) but induce *pilA9-pilA12* transcript accumulation, which challenges our simplified model. Another mRNA encoding the minor pilin *pilX1* (*slr0226*), which is involved in cellular aggregation, responds to high c-di-GMP levels (Wallner et al., 2020; Oeser et al., 2021) and is downregulated in the *cdgR* mutant (Fig. 4A). Interestingly, *pilX1* mutant cells do not flocculate, but are motile and transformable (Oeser et al., 2021). *Synechocystis* encodes 13 different minor pilins which are involved in the different functions of type IV pili and are most probably part of distinct pilus fibers. Therefore, we speculate that the ratio between different minor pilins defines the number of specific pilus fibers, eventually leading to pleiotropic phenotypes. Moreover, cAMP is an important, yet not fully understood, player in motility control in cyanobacteria. This second messenger is known to activate SyCRP1, and *cyaA* mutants are not motile, suggesting that cyanobacterial type IV pili may be involved in surface-induced cAMP signaling, as described in *Pseudomonas aeruginosa* (Persat et al., 2015; Kühn et al., 2021).

**Figure 7:**
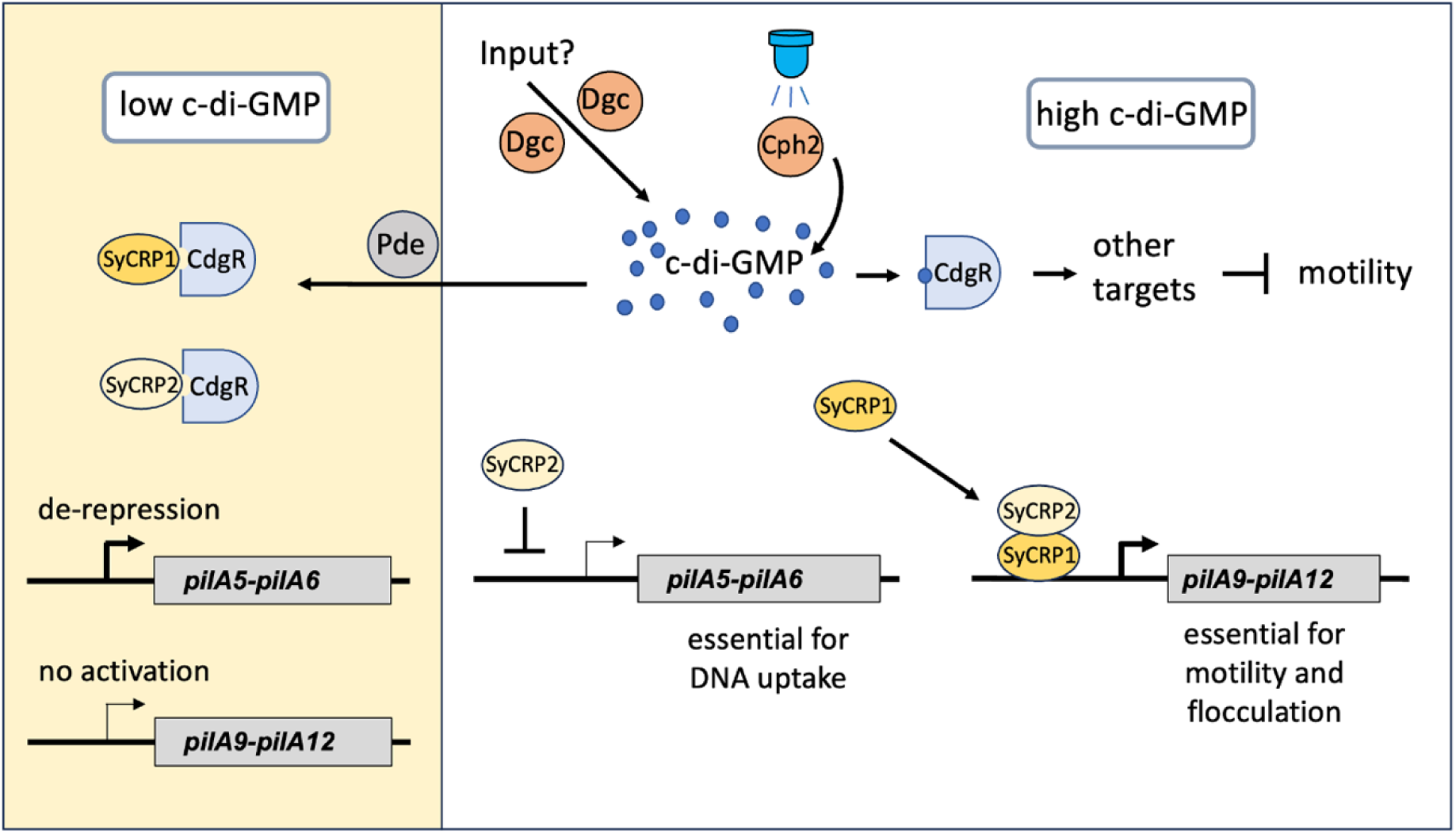
Model of the function of CdgR in the control of c-di-GMP-dependent expression of minor pilins. Under high c-di-GMP conditions (e.g., blue light-dependent cyclase activity of Cph2), CdgR does not bind to the transcription factors SyCRP1 and SyCRP2. Therefore, they can bind to DNA and repress or activate the expression of minor pilin genes. Under low cellular c-di-GMP concentrations, CdgR inactivates both transcription factors by binding to them. In the absence of CdgR, the *pilA5-pilA6* operon, which encodes minor pilins important for DNA uptake, is repressed, and the cells are not transformable. Overaccumulation of *pilA9-pilA12* mRNA in the *cdgR* mutant leads to enhanced phototactic movement. Other unknown functions of CdgR beyond the control of the two CRP-like transcription factors, as well as the contribution of other nucleotide second messengers, are likely.

In contrast to the multicellular cyanobacterium *Anabaena*, where *cdgR* controls cell size, *cdgR* (in *Synechocystis*) or its homolog (in *Bacillus* or *Shewanella*) (Hahn et al., 2025; Samir et al., 2025b) appears to have a conserved function in controlling bacterial motility in a c-di-GMP-dependent manner. However, the mechanism by which *Bacillus* or *Shewanella* homologs control motility remains unclear (Hahn et al., 2025; Samir et al., 2025b). Here, we showed that CdgR binds more specifically to c-di-GMP, but c-di-AMP can substitute for c-di-GMP at higher concentrations. This raises the question of whether c-di-AMP can replace c-di-GMP *in vivo* under conditions where c-di-GMP levels are low and c-di-AMP is high. In *Synechocystis*, the c-di-GMP concentration was one order of magnitude higher than that of c-di-AMP under constant white light conditions (Supplementary Figure S4). This aligns with our previous finding that c-di-GMP levels are 10 times higher than c-di-AMP levels under both day and night conditions (Selim et al., 2021; Samir et al., 2025a). Therefore, we assume that under our *in vivo* conditions, CdgR preferentially binds c-di-GMP under our *in vivo* conditions. However, other conditions, such as osmotic stress, could increase c-di-AMP levels (Agostoni et al., 2018), and consequently change the c-di-GMP/c-di-AMP ratio.

Crosstalk between second messenger nucleotides, as demonstrated for CdgR, may be a more common phenomenon than previously realized. This is observed, for instance, between different cyclic di-nucleotides and other messengers. In *Caulobacter crescentus*, c-di-GMP and (p)ppGpp reciprocally control growth by competitively binding to the metabolic switch protein SmbA (Shyp et al., 2021). Similarly, crosstalk between c-di-AMP and cAMP is mediated by SbtB in cyanobacteria (Selim et al., 2018; Forchhammer et al., 2022; Selim and Alva, 2024) and the transcription factor DarR in mycobacteria (Schumacher et al., 2023), which bind both molecules. Crosstalk is also well documented between cAMP and cGMP. For example, CRP-Fnr transcription factors bind both signaling nucleotides. However, their activation differs by species; only the cAMP-bound form is active in *E. coli*, whereas both nucleotides activate CRP in *Sinorhizobium meliloti* (Krol et al., 2023; Werel et al., 2023).

Surprisingly, we found that the CdgR-SyCRP1 system integrates both cAMP and c-di-NMP signals, with SyCRP1 binding cAMP and CdgR binding c-di-NMPs. The binding of cAMP to SyCRP1 had a protective effect, preventing the dissociation of the CdgR-SyCRP1 complex. This was not the case for SyCRP2, which does not bind to cAMP. Such crosstalk between different second messengers, mediated by two distinct proteins, has recently been recognized. For instance, the activity of (p)ppGpp synthetase in *Bacillus* is controlled by its interaction with the c-di-AMP receptor protein DarB in a c-di-AMP-dependent manner, which consequently controls the cellular levels of (p)ppGpp (Krüger et al., 2021).

Our results on the *ΔcdgR/Δcph2* double mutant suggest that the blue-light-dependent c-di-GMP signaling pathway which controls motility in *Synechocystis* might also operate independently of CdgR. This interpretation is consistent with the growing evidence from other bacteria that c-di-GMP signaling can be highly compartmentalized, with distinct receptor proteins responding to specific local c-di-GMP pools rather than global intracellular concentrations. Therefore, we assume that, beyond CdgR, there are more potential c-di-GMP-dependent effectors in *Synechocystis* which can independently regulate type IV pilus functions. Five potential c-di-GMP-binding proteins with an MshEN domain (including the type IV pilus assembly ATPase PilB) have been identified in *Synechocystis* (Wang et al., 2016). Indeed, it has been shown that PilB can bind c-di-GMP in heterotrophic bacteria, and that this interaction is important for the switch between motility and biofilm formation in *Myxococcus xanthus* (Dye et al., 2023). Identification of additional c-di-GMP receptors and their relationship to Cph2-derived signaling will therefore be important for understanding how light-specific motility responses are integrated in cyanobacteria. As photosynthetic organisms, cyanobacteria essentially depend on light availability. Type IV pili-dependent motility allows cyanobacteria to move toward optimal light qualities and intensities or escape dangerous light conditions by negative phototaxis or aggregation. When cyanobacteria aggregate, whether by forming dense clumps, multi-layered mats, or biofilms, they transition from vulnerable, isolated individuals into a cooperative, structured community. This collective physical structure combined with the induction of DNA uptake and recombination, is effective at neutralizing the dangers of high-intensity solar radiation, including UV light. Thus, cyanobacteria are able to suppress motility and induce aggregation and/or DNA uptake when it is necessary to escape from stressful situations.

In summary, a more complex model beyond the current CdgR-centered model, including cross-talk between different second messengers, multiple signals influencing their concentrations, and several effectors which respond to different concentrations and ratios of second messengers, needs to be developed to fully understand how c-di-GMP controls type IV pilus-dependent functions.

## Acknowledgements

This project was funded by the National Natural Science Foundation of China (Grant Nos. 32170055) to Y.Y. and DFG as part of SFB1381 (project number: 403222702) to K.A.S. and A.W and Emmy Noether program (SE 3449/3-1) to K.A.S. We thank Karl Forchhammer for his continued support and constructive discussions. We acknowledge the infrastructural support from CMFI (EXC 2124-390838134) and SFB1535 MibiNet (project number: 458090666). S.S. was supported by the Egyptian Ministry of Higher Education, Egypt. We thank Filipp Oesterhelt (Tübingen University), John Weir (MPI for Biology; Tübingen) and Daniel Wohlwend (Freiburg University) for facilitating the usage of mass photometry or ITC. We thank Zhi-xian Qiao at the Analysis and Testing Center of Institute of Hydrobiology, Chinese Academy of Sciences for her assistance in RNAseq analysis.

## Author contributions

A.W., C.Z., Y.Y. and K.A.S. conceived the project; T.W., Y.Y. and K.A.S. designed research; C.H., T.W., S.S., X.Z and E.S.L. performed research; A.W., C.Z. and K.A.S. contributed reagents/analytic tools; Y.Y., T.W., C.H., S.S., and K.A.S. analyzed data; and T.W., A.W., Y.Y. and K.A.S. wrote the paper.

## Data availability statement

The full data set derived from the RNAseq analysis is accessible in the Gene Expression Omnibus database with the accession number GSE322604 (https://www.ncbi.nlm.nih.gov/geo/query/acc.cgi?acc=GSE322604; security access token for reviewers: cxkpgyyavjshfyv)

## Statement on generative AI and AI-assisted technologies in the writing process

The authors used AI-assisted technologies such as paperpal.com and neural machine translation such as DeepL.com to improve readability and correct linguistic errors. After using these tools/services, the authors manually reviewed and edited the content. The authors take full responsibility for the content of the published article.

