## supplemental material for "Regulation of cyanobacterial type IV pilus-dependent functions by interaction between a c-di-GMP receptor and two transcription factors"

**The pdf file includes:**

Supplementary figures S1-S11

Supplementary tables S1-S3

Supplementary references

**Other Supplementary Material for this manuscript (not in this pdf) include:**

Supplementary table S4 (excel file): RNA-seq dataset of ∆*cdgR* mutant compared to the wild type

Supplementary table S5 (excel file): statistical analysis of phototactic movement

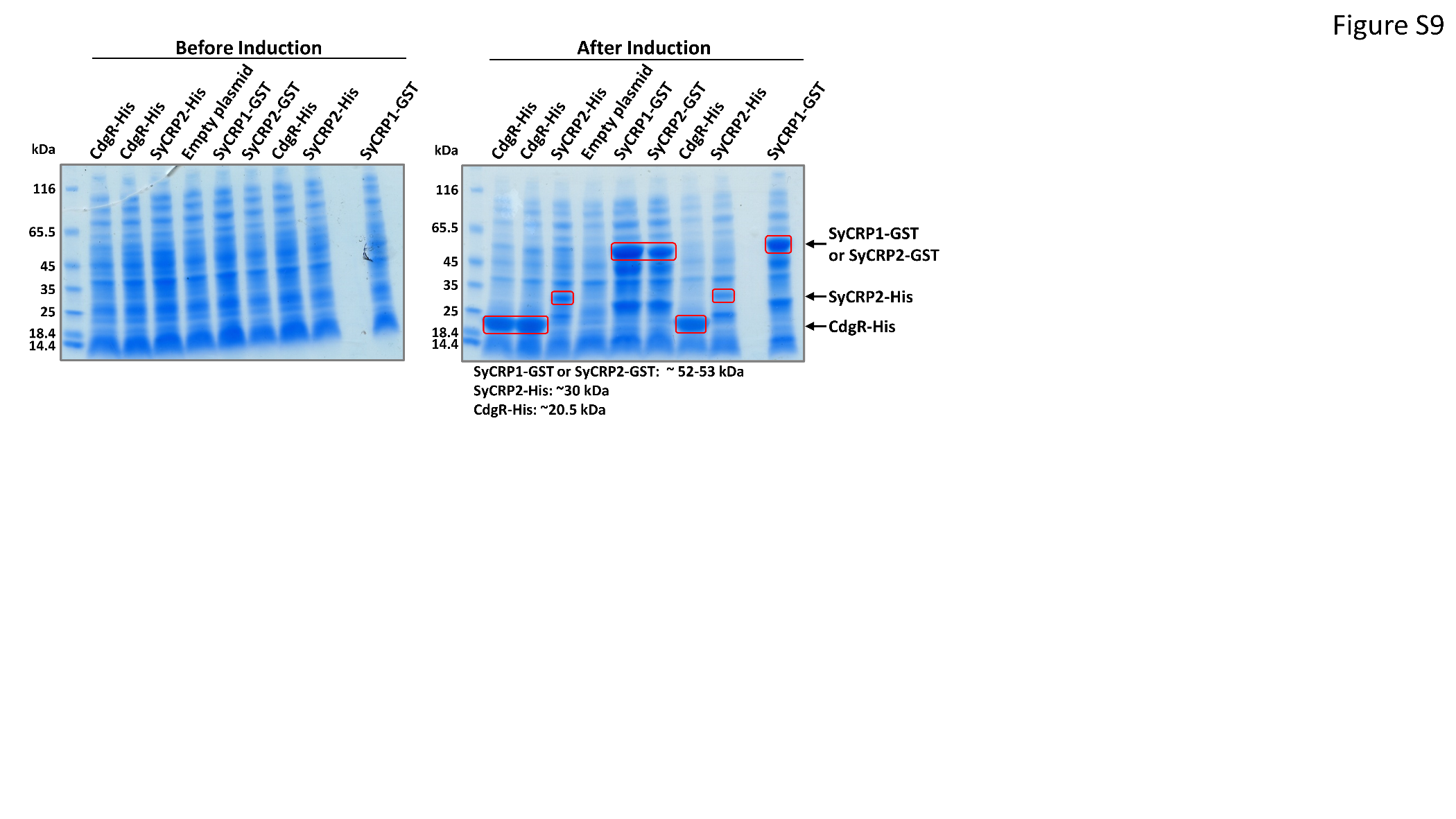

**Supplementary Figure S1: SDS-PAGE analysis of tagged protein expression.** Crude-cell lysates from *E. coli* BL21(DE3) cells expressing CdgR-His, SyCRP2-His, SyCRP1-GST, or SyCRP2-GST proteins were separated before and after induction. Proteins were visualized by Coomassie staining. Target bands are outlined in red boxes, and molecular weights are indicated.

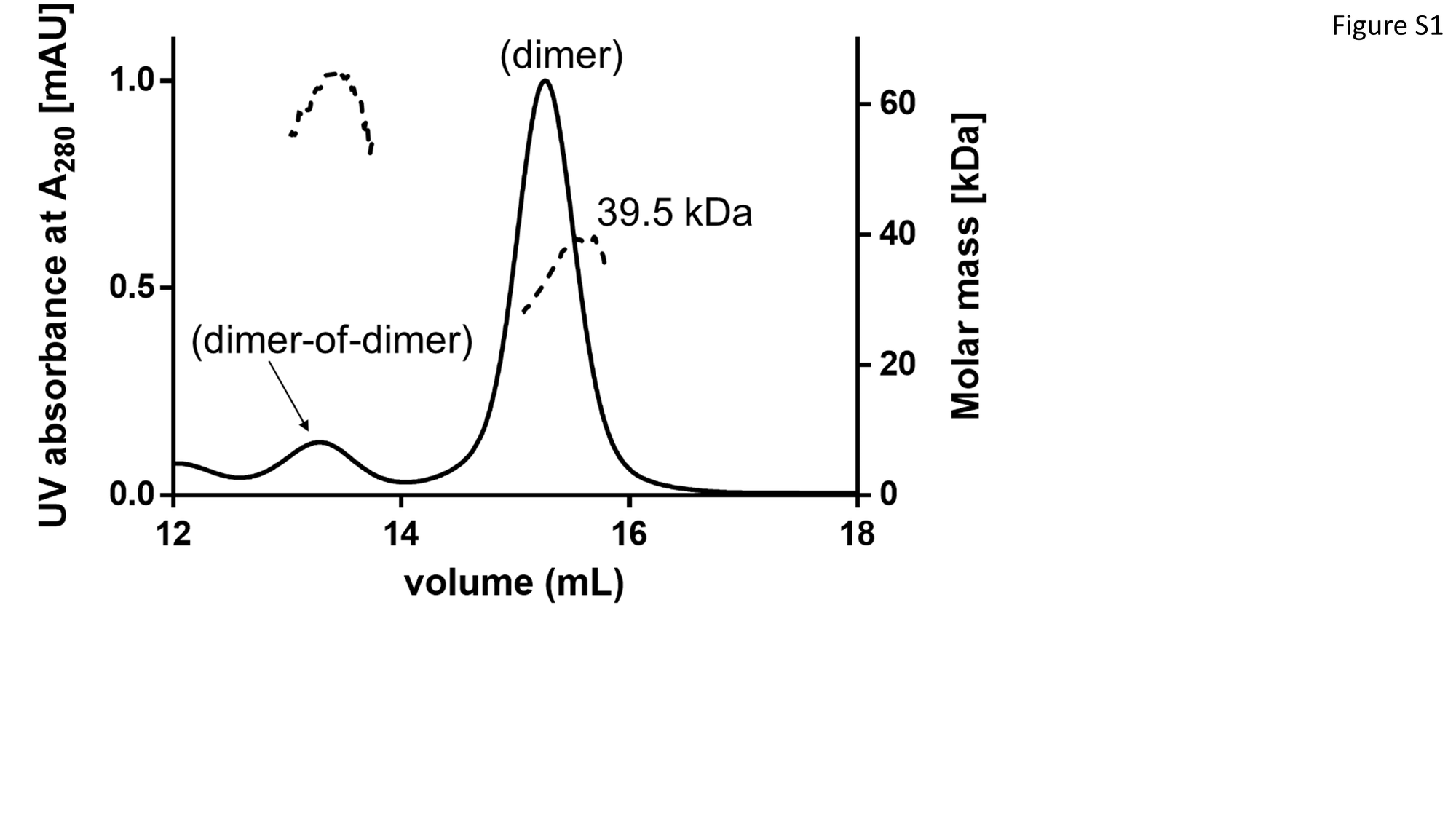

**Supplementary Figure S2: Purification of native *Synechocystis* CdgR and its oligomeric state.** Analytical size-exclusion chromatography coupled to multi-angle light scattering (SEC-MALS) was conducted as described previously to calculate the molar mass of the CdgR protein (Samir et al., 2025a).

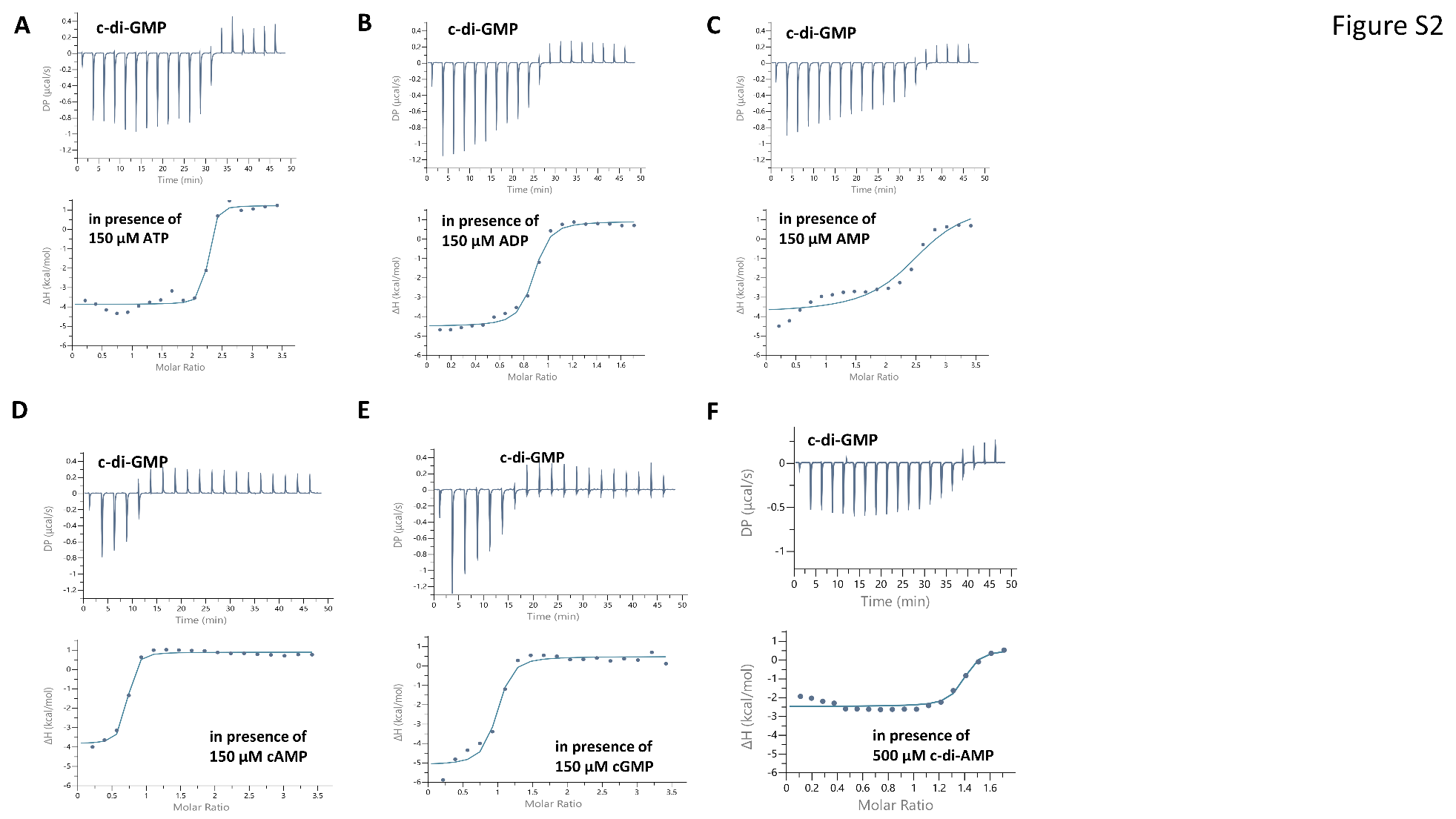

**Supplementary Figure S3: ITC analysis of c-di-GMP binding to CdgR in the presence of the major cellular nucleotides or other cyclic second messengers.** Upper panels show the raw ITC data in the form of heat produced during the titration of c-di-GMP in presence of 150 µM of **(A)** ATP, **(B)** ADP, **(C)** AMP, **(D)** cAMP, **(E)** cGMP, and **(F)** 500 µM c-di-AMP. Lower panels show the binding isotherms and the best-fit curves according to the one binding site model.

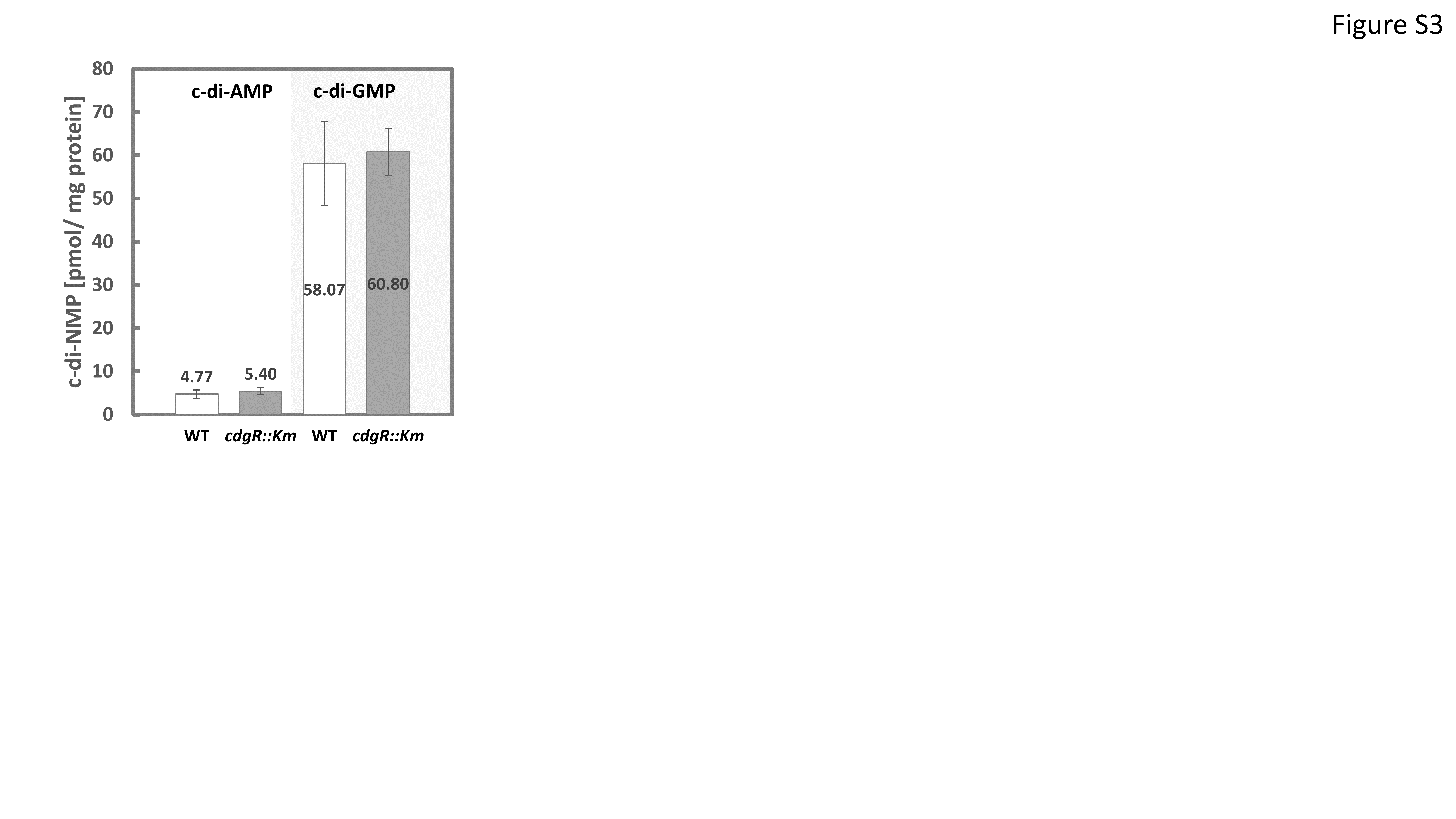

**Supplementary Figure S4: Cellular levels of cyclic nucleotide messengers in the *cdgR::Km* mutant strain.** Cells of *Synechocystis* wild type and the *cdgR::Km* mutant were grown in BG11 medium with 75 µmol photons m^-2^ s^-1^ white-light illumination, and the content of the cyclic nucleotide second messengers c-di-AMP (left) and c-di-GMP (right) was determined in the logarithmic growth phase. The content of each second messenger was normalized to the total protein content of the cells. Two or four biological replicates with technical triplicates were used. Wild type, WT.

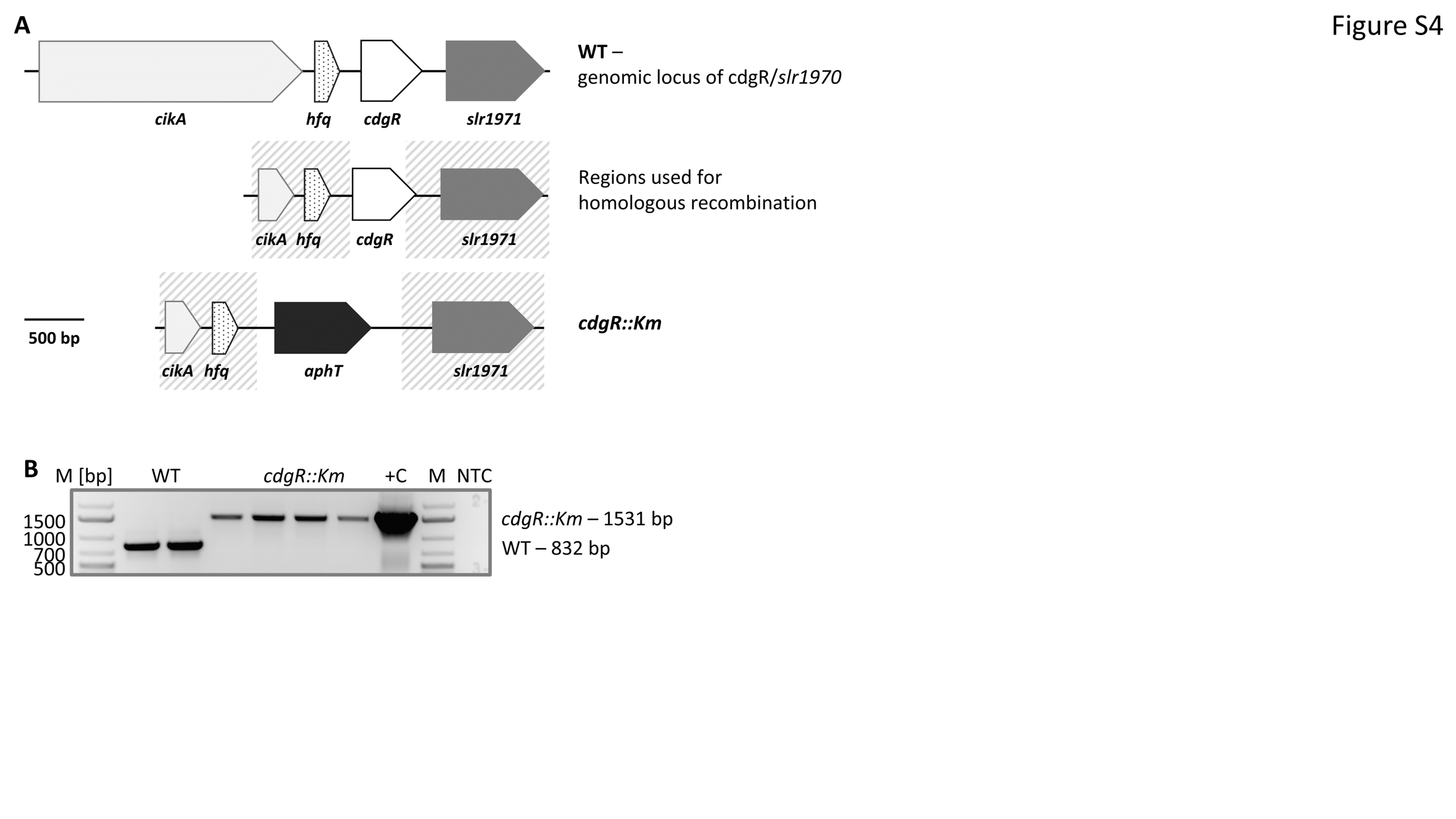

**Supplementary Figure S5: Schematic representation of the *cdgR* genomic context and verification of the deletion of the *cdgR* gene. A:** The gene locus of *cdgR/slr1970* is displayed*,* along with the co-transcribed gene *hfq/ssr3341* (transcriptional unit TU1845 according to Kopf et al. (2014)), the upstream gene *cikA/slr1969,* and the downstream gene *slr1971.* The gray stripe boxes indicate the regions utilized for homologous recombination with the *Synechocystis* chromosome. To completely replace *cdgR*´s entire sequence, except for the last 33 bp, a kanamycin resistance gene cassette (*aphT*) was used. **B:** Detection of the complete segregation of mutant copies of *cdgR::Km* was achieved using colony PCR and the primer pair *slr1970*-col-fw and *slr1970*-col-rev (Table S1). Negative controls were chromosomal wild-type DNA and a non-template reaction (**NTC**), while the plasmid used for transforming *Synechocystis* wild type was the positive control (**+C**). The expected sizes for the wild type and the *cdgR::Km* allele are 832 bp and 1531 bp, respectively. **M [bp]**, DNA molecular weight standard.

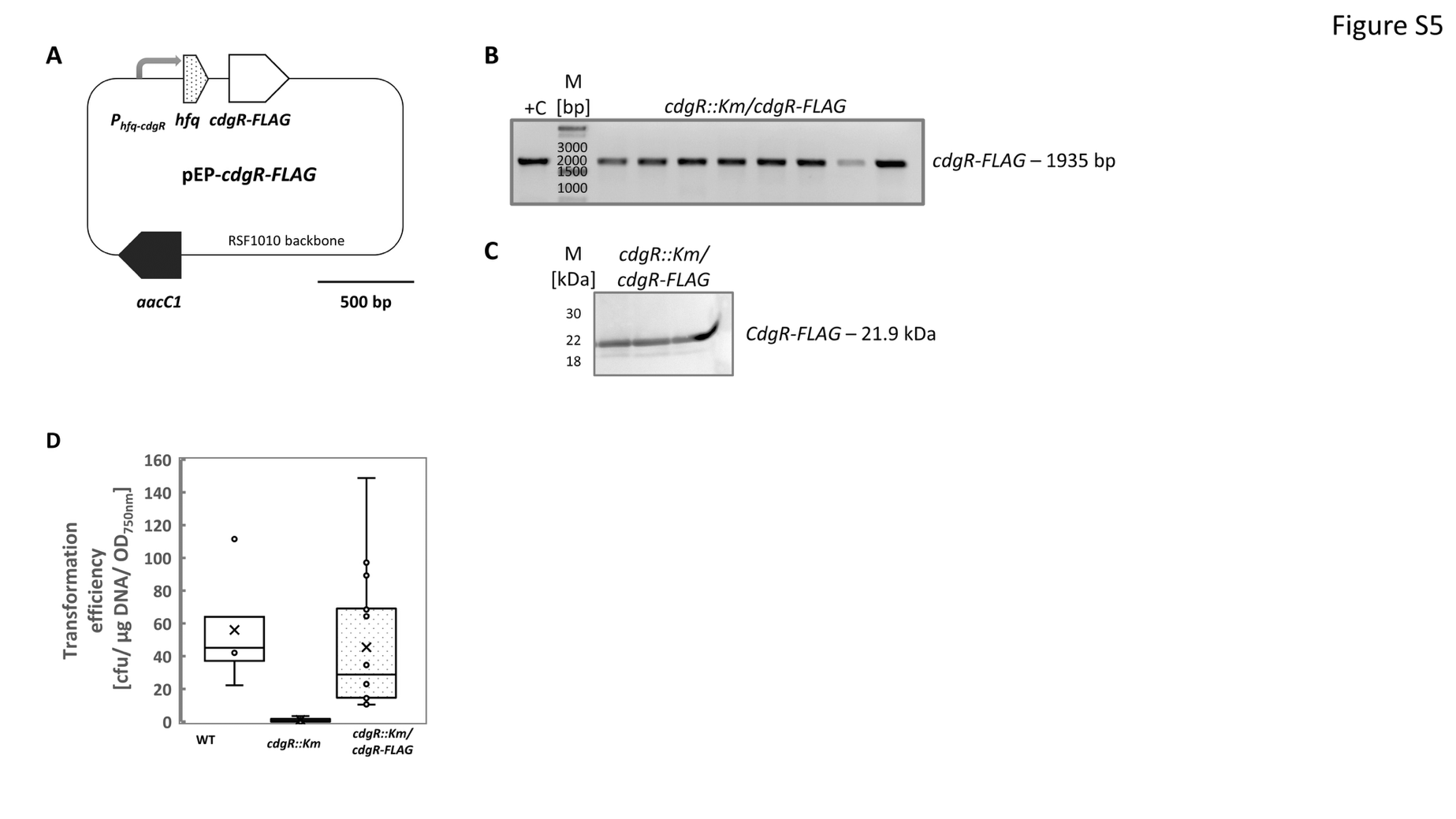

**Supplementary Fig. S6: Schematic representation of the complementation of the *cdgR::Km* mutant. A:** The entire TU1845 transcription unit (Kopf et al., 2014), consisting of *hfq/ssr3441* and *cdgR/slr1970,* was amplified from *Synechocystis* genomic DNA and ligated into the pJET vector. This amplicon contained the native promoter of TU1845. The NheI restriction enzyme recognition sequence was introduced in-frame at alanine 169 position of CdgR using the Q5 site-directed mutagenesis kit (New England Biolabs). A fused oligonucleotide encoding a triple FLAG tag and a stop codon was ligated into the NheI site. The resulting construct was excised from the pJET vector using HindIII and BamHI and ligated into a modified conjugative pVZ322 plasmid. **B:** Detection of the pEP-*cdgR*-FLAG plasmid in the *cdgR::Km*/*cdgR*-FLAG complementation strain was achieved by colony PCR using the primer pair *aaC1*-seq-out-fw and pUR-rev (Table S1). The plasmid used for conjugation of the *cdgR::Km* mutant strain served as a positive control (**+C**). The expected size is 1935 bp. **M [bp]**, DNA molecular weight standard. **C:** Detection of CdgR-FLAG protein in the *cdgR::Km*/*cdgR*-FLAG complementation strain. Cells were scraped from agar plates, resuspended in SDS loading buffer (Laemmli, 1970), and total protein was denatured at 50 °C for 30 min. Proteins were separated on a 4-12% gradient gel and blotted onto a nitrocellulose membrane. The CdgR-FLAG fusion protein was detected after incubation with horseradish peroxidase-conjugated α-FLAG M2 antibody (Sigma-Aldrich). The expected size of the target protein was 21.9 kDa. **M [kDa]**, protein molecular weight standard. **D:** Transformation efficiency of the *cdgR::Km/cdgR*-FLAG complementation strain. These experiments determined the transformation efficiency of *Synechocystis* wild type (n = 4) and the *cdgR::Km* mutant strain (n = 4), as well as of the *cdgR::Km/cdgR*-FLAG complementation strain (n=16). The number of colony-forming units (cfu) was counted after 12 days, and the transformation efficiency was normalized to the optical density at 750 nm (OD_750nm_) and the amount of DNA used. For each transformation, one microgram of pJET*-ΔcrhR*-SpecR plasmid DNA was used, conferring spectinomycin resistance.

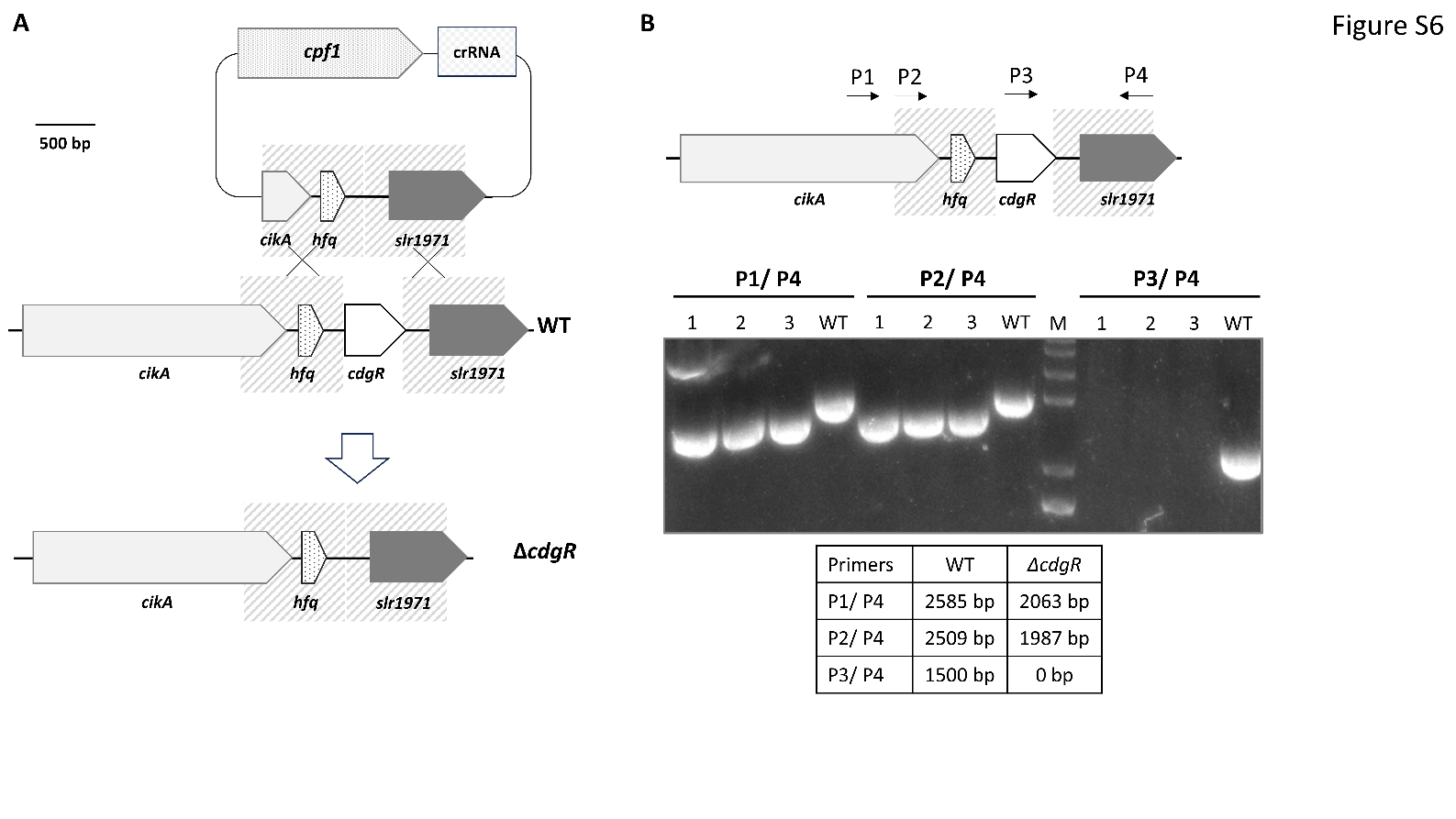

**Supplementary Figure S7: CRISPR-Cas12a (Cpf1)-mediated deletion of the *cdgR* gene. A**: Strategy for constructing the gene knockout plasmid. A crRNA sequence targeting the *cdgR* locus (*slr1970*) and homologous flanking sequences (shaded area) were cloned into the pCpf1-sp vector to generate the donor plasmid pCpf1-*cdgR-*sp, designed for in-frame deletion of the *cdgR* gene. **B**: PCR genotyping of mutant candidates. Three independent transformants (clone #1-3) and a wild-type control were verified using primer pairs as indicated. The sizes of the expected PCR products for the wild-type and mutant alleles are summarized in the table.

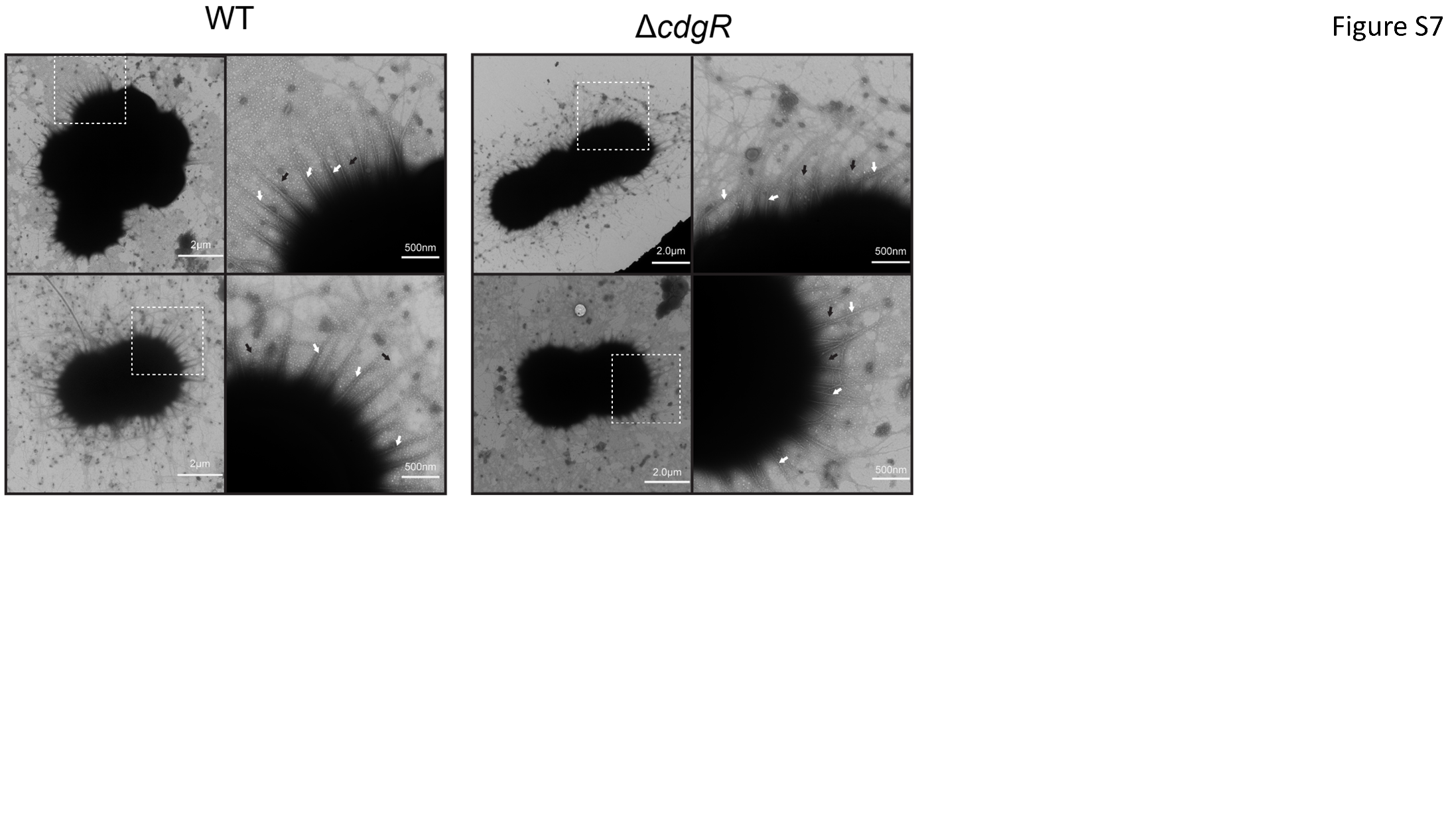

**Supplementary Figure S8: Electron micrographs of negatively stained WT and *ΔcdgR* deletion mutants.** Whole-cell images and magnified views of the dashed areas are shown. White arrows indicate thick pili, and black arrows indicate thin pili (Bhaya et al., 2000; Yoshihara et al., 2001).

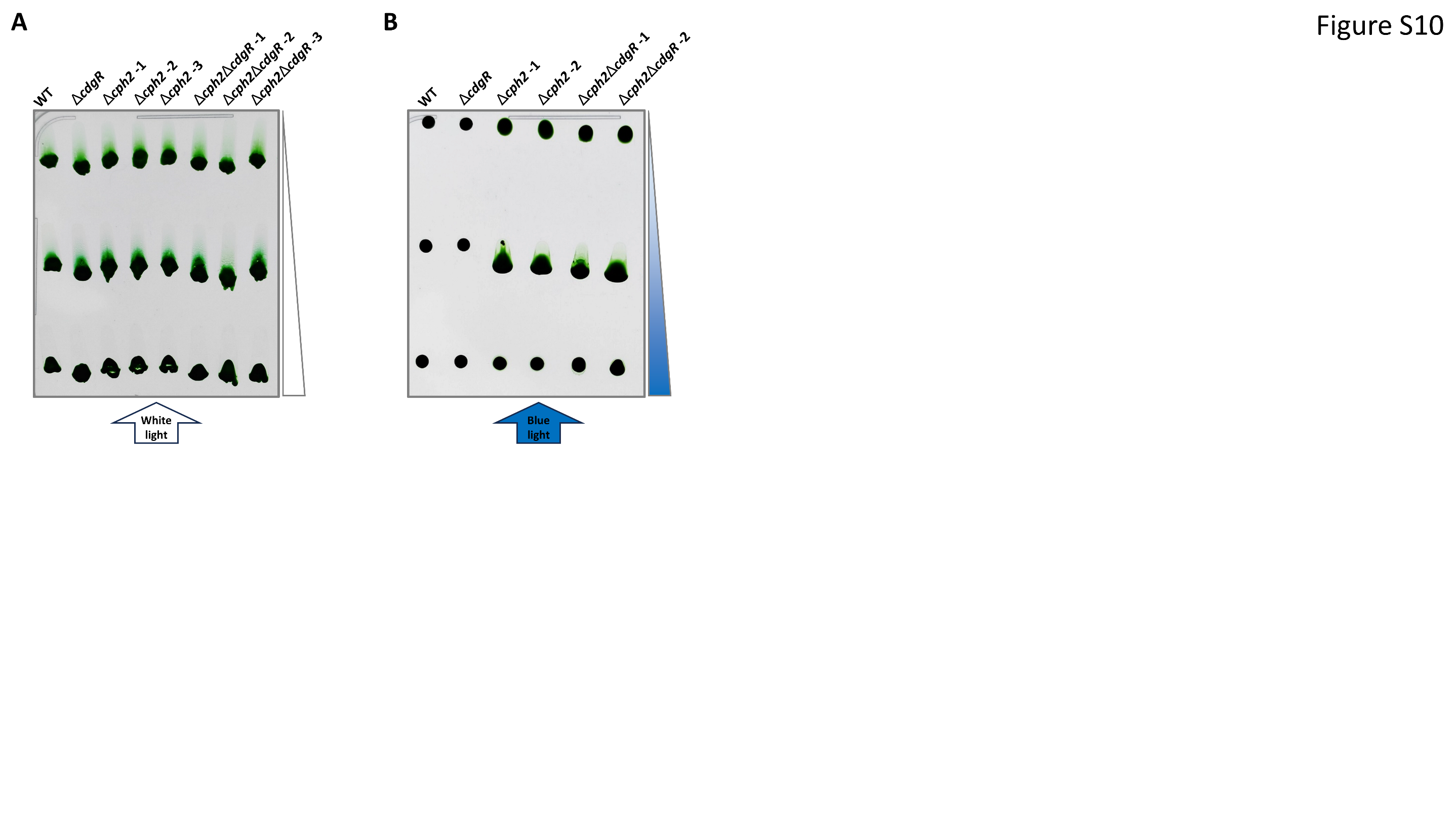

**Supplementary Figure S9. Phototactic motility of *cdgR* and *cph2* single and double mutants.** Motility assays were performed on BG11 plates (0.7% agar) supplemented with 2 mM TP and 3.3 mM Cu^2+^. Two to three independent mutant clones of Δ*cph2* and Δ*cph2/*Δ*cdgR* were incubated under lateral illumination with white light for 6 days **(A)** or blue light for 12 days **(B)**. Light intensity gradients were indicated by the white and blue triangles.

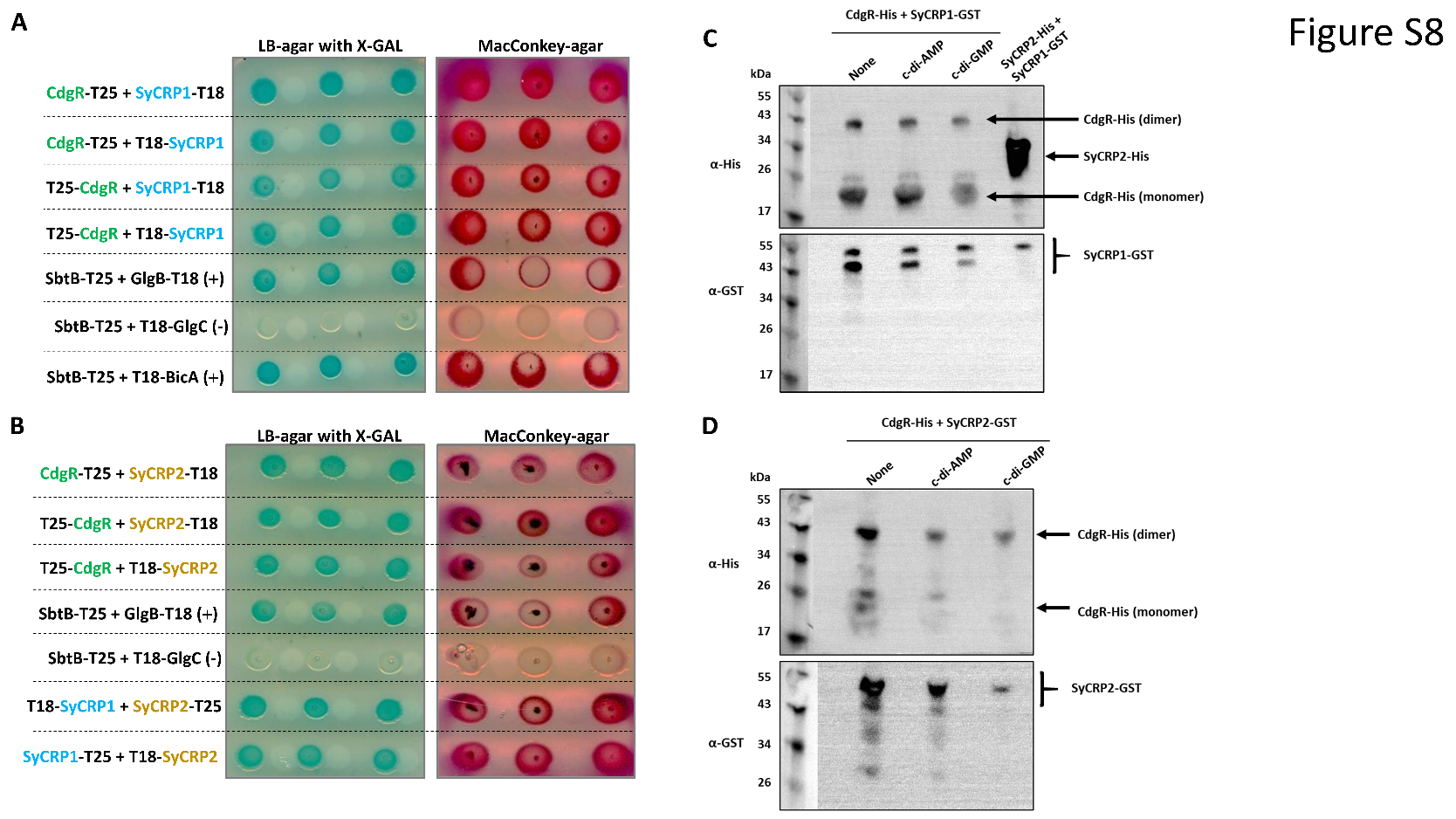

**Supplementary Figure S10: Screening for CdgR-interacting proteins.**

**(A, B):** The BACTH assay was employed to identify proteins that interact with CdgR. Strains of *E. coli* BTH101 cells co-expressing CdgR and potentially interacting proteins fused to the T18 or T25-fragment of the *Bordetella pertussis* adenylate cyclase CyaA were applied onto MacConkey-agar plates containing 1% (w/v) maltose and 1 mM IPTG or onto LB-agar plates containing X-GAL (40 µg/ml) and IPTG (1 mM). The plates were then incubated at 25°C for up to 96 h. The *Synechocystis* carbon control protein SbtB (*slr1513*), glycogen-branching enzyme (GlgB, *sll0158*), and bicarbonate transport BicA (*sll0834*) served as positive controls (**+**), while glucose-1-phosphate adenylyltransferase (GlgC, *slr1176*) acted as a negative control (Selim et al., 2021; Haffner et al., 2025). Three individual clones were procured from the co-transformations. **A:** *Synechocystis* 6803 cAMP receptor protein SyCRP1, and **B:** *Synechocystis* 6803 cAMP receptor like protein SyCRP2.

**(C, D): Immunoblot blot analysis of CdgR and SyCRP1/2 interaction by in vitro pulldown assays.** In the pulldown assays, the binding between CdgR-His (immobilized on Ni²⁺-NTA beads) and SyCRP1-GST (C) or SyCRP2-GST (D) (in the crude extract) was assessed by Western blot using α-His or α-GST antibodies, respectively. Assays were conducted in the absence (None) or presence of 0.5 mM c-di-AMP or c-di-GMP to investigate their potential modulatory roles. Proteins were detected via Western blotting using anti-His (for CdgR) and anti-GST (for SyCRP1/2) antibodies. **Top panels:** anti-His western blot for detecting CdgR (monomer: 20.5 kDa; dimer: ~43 kDa). **Bottom panels:** anti-GST Western blot for detecting SyCRP1-GST (C; 52.2 kDa) or SyCRP2-GST (D; 53.3 KDa) fusion proteins. The smaller signal at about 43 kDa with the GST antibody derives from partial degradation of the GST-tagged transcription factors. Lane 4 in panel (C): positive control of SyCRP2-His (~30 kDa) and SyCRP1-GST (~52.2 kDa) interaction.

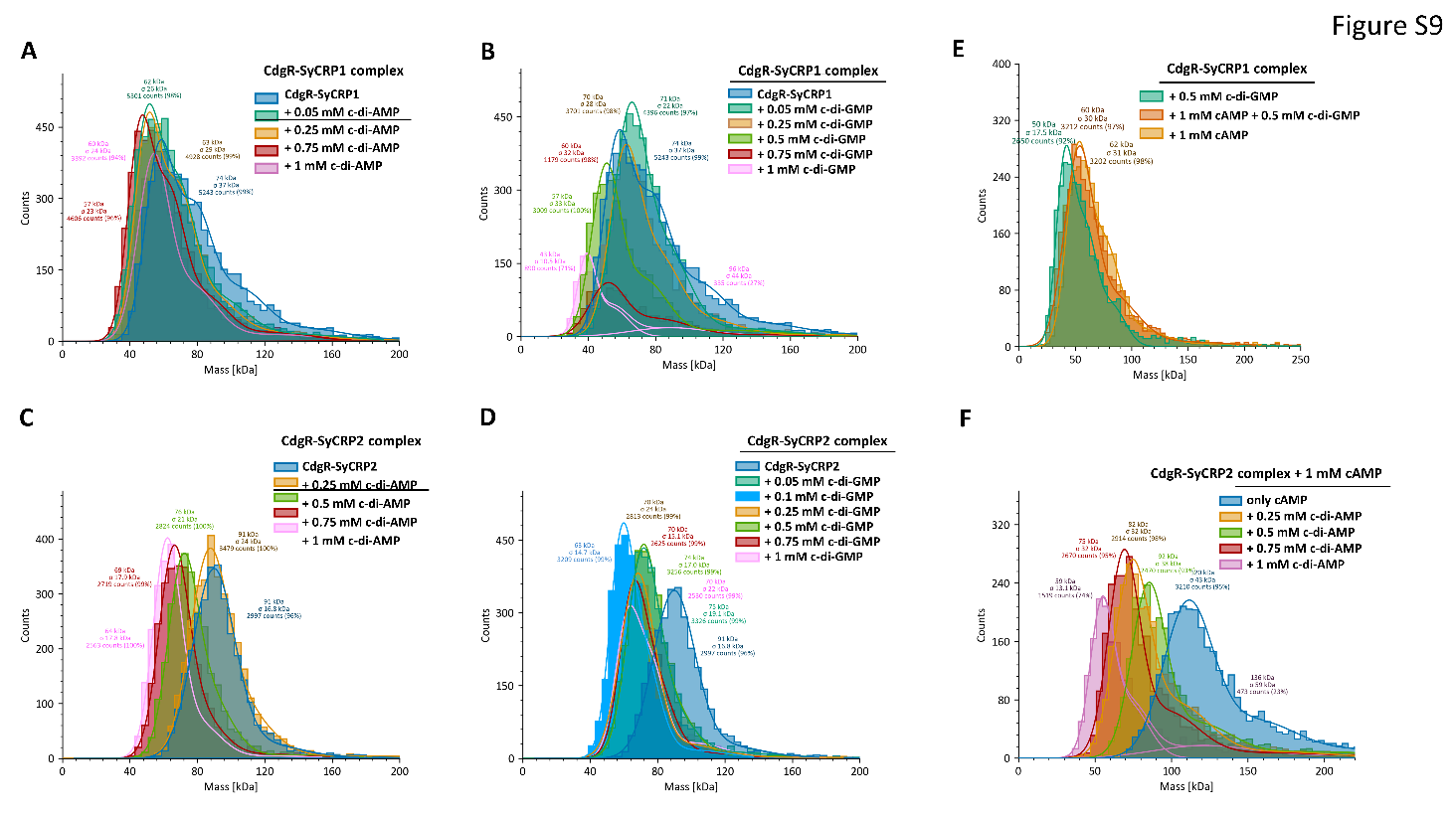

**Supplementary Figure S11: Mass photometry showing the CdgR-SyCRP1/2 complex formation.** The effects of increasing concentrations of c-di-AMP **(A)** or c-di-GMP **(B)** on CdgR-SyCPR1 complex formation is shown. The effects of increasing concentrations of c-di-AMP **(C)** or c-di-GMP **(D)** on CdgR-SyCRP2 complex formation are depicted. The effect of cAMP on CdgR-SyCRP1 complex in absence and presence of c-di-GMP is shown in **(E)**. The effects of increasing concentrations of c-di-AMP on the CdgR-SyCRP2 complex in the presence of cAMP is shown in panel **F**. The complexes are indicated by higher mass of 90-120 kDa.

**Supplementary Table S1.** Primers used in this study.

| **Name** | **Sequence (5´-3´)** | **purpose** |
| --- | --- | --- |
| AQ-*slr1970*-P1 | CTTAACTATGCGGCATCGGTCATTTGGTTAATTG | Construction of pUC19-*cdgR*-Km |
| AQ-*slr1970*-P2 | TGAGACACAACGTGGCGGTAGAAAGGCTGATTGG | Construction of pUC19-*cdgR*-Km |
| AQ-*slr1970*-P3 | CCAATCAGCCTTTCTACCGCCACGTTGTGTCTCA | Construction of pUC19-*cdgR*-Km |
| AQ-*slr1970*-P4 | GCTTCCAATAACAATGCGAGGTCTGCCTCGTGAA | Construction of pUC19-*cdgR*-Km |
| AQ-*slr1970*-P5 | TTCACGAGGCAGACCTCGCATTGTTATTGGAAGC | Construction of pUC19-*cdgR*-Km |
| AQ-*slr1970*-P6 | CACTTTATGCTTCCGGCCCACCGTCACTTTCACT | Construction of pUC19-*cdgR*-Km |
| AQ-*slr1970*-P7 | AGTGAAAGTGACGGTGGGCCGGAAGCATAAAGTG | Construction of pUC19-*cdgR*-Km |
| AQ-*slr1970*-P8 | CAATTAACCAAATGACCGATGCCGCATAGTTAAG | Construction of pUC19-*cdgR*-Km |
| *slr1970*-col-fw | CCAGTCAAAGCAGTTATAGC | Segregation verification of *cdgR*::Km |
| *slr1970*-col-rev | GAACCAACCAGGGTAAAAC | Segregation verification of *cdgR*::Km |
| 1970_C-FW | GCTAAGCTTGGTCATTTGGTTAATTGATACTACG | Construction of pEP-*cdgR*-FLAG |
| 1970_C-REV | GCTGGATCCCACTAGGCCCAATTAATCCAC | Construction of pEP-*cdgR*-FLAG |
| A168A-NheI-FW | TATTGGAAGCGCTAGCCCAGG | Construction of pEP-*cdgR*-FLAG |
| A168A-NheI-REV | ACAATGCCATTTGACTGATCAG | Construction of pEP-*cdgR*-FLAG |
| XbaI-FLAG-fw | CTAGATTATAAAGATCATGATGGCGATTATAAAGATCATGATATTGATTATAAAGATGATGATGATAAAGG | Construction of pEP-*cdgR*-FLAG |
| XbaI-FLAG-reverse | CTAGCCTTTATCATCATCATCTTTATAATCAATATCATGATCTTTATAATCGCCATCATGATCTTTATAAT | Construction of pEP-*cdgR*-FLAG |
| aaC1-seq-out-fw | CGGTAAATTGTCACAACGCC | Detection of pEP-*cdgR*-FLAG |
| pUR-rev | CTTCCAGATGTATGCTCTTCTGCTC | Detection of pEP-*cdgR*-FLAG |
| slr1928-pilA5-fw | CCCCAGGCTTTTATCTCTCG | Generation of a T7 DNA template for a probe targeting *pilA5* mRNA |
| T7-slr1928-pilA5-rev | **TAATACGACTCACTATAGGG**GGCGTCAGCACTATTGTTTG | Generation of a T7 DNA template for a probe targeting *pilA5* mRNA |
| pilA9-fw | CTAGCGACCGATACCACCAT | Generation of a T7 DNA template for a probe targeting *pilA9* mRNA |
| T7-pilA9-rev | **TAATACGACTCACTATAGGG**ACGCCGTTATTGCATTTTTC | Generation of a T7 DNA template for a probe targeting *pilA9* mRNA |
| 16S_fw | CAAGTCATCATGCCCCTTAC | generation of dsDNA probe targeting 16S rRNA |
| 16S_rev | GATGGGATTCGCTTACTCTC | Generation of dsDNA probe targeting 16S rRNA |
| BACTH-Slr1970-fw | GA**GGATCC**GAGCATCCAAGAAATCTTTACC | Construction of BACTH plasmids pUT18/pUT18C/pKT25 and pKNT25 expressing CdgR/Slr1970 |
| 25/18-Slr1970-rev | GT**GGTACC**ATCCTGGGCTAGGGC | Construction of BACTH plasmids pUT18/pUT18C/pKT25 and pKNT25 expressing CdgR/Slr1970 |
| Slr1970-25/18-rev | GT**GGTACCTT**ATCCTGGGCTAGGGC | Construction of BACTH plasmids pUT18/pUT18C/pKT25 and pKNT25 expressing CdgR/Slr1970 |
| BACTH-SyCRP1-fw | CA**GGATCC**AGGCACTAGTCCCCAAAATTC | Construction of BACTH plasmids pUT18/pUT18C/pKT25 and pKNT25 expressing SyCRP1/Sll1371 |
| 25/18-SyCRP1-rev | GT**GGTACC**GGAAATTAGATCTTCTAAATCCC | Construction of BACTH plasmids pUT18/pUT18C/pKT25 and pKNT25 expressing SyCRP1/Sll1371 |
| SyCRP1-25/18-rev | GT**GGTACC**GTGGAAATTAGATCTTCTAAATCCC | Construction of BACTH plasmids pUT18/pUT18C/pKT25 and pKNT25 expressing SyCRP1/Sll1371 |
| BACTH-sycrp2-fw | CA**GGATCC**CGCACCACAAAAGCCGTCAAC | Construction of BACTH plasmids pUT18/pUT18C/pKT25 and pKNT25 expressing SyCRP2/Sll1924 |
| 25/18-sycrp2-rev | CA**GGTACC**GATCGAAGTGGGATTGCCAC | Construction of BACTH plasmids pUT18/pUT18C/pKT25 and pKNT25 expressing SyCRP2/Sll1924 |
| sycrp2-25/18-rev | CA**GGTACCGG**GATCGAAGTGGGATTGCCAC | Construction of BACTH plasmids pUT18/pUT18C/pKT25 and pKNT25 expressing SyCRP2/Sll1924 |
| PV_14 | GCAATGGCAACAACGTTGCG | Linearization of pCpf1b-sp vector |
| p*sacB*R | GATTTGCAGCATATCATGGCGTGT | Linearization of pCpf1b-sp vector |
| p*sacB*F | ACACGCCATGATATGCTGCAAATC | Linearization of pCpf1b-sp vector |
| Pcpf1R3795 | CCTACCTAGTAGCATCAGACCT | Linearization of pCpf1b-sp vector |
| Pcpf1F3795 | AGGTCTGATGCTACTAGGTAGG | Amplification of *slr1970* from pCpf1-Mslr1970R236(180302F22) |
| P*slr1970*R2510 | CGCAACGTTGTTGCCATTGTCCGTCTCCTTGGTAGAGGTA | Amplification of *slr1970* from pCpf1-M*slr1970*R236(180302F22) |
| pUC19-2 | AAGCTTGGCGTAATCATGGTC | Amplification of pUC19 |
| pUC19-1 | GAATTCACTGGCCGTCGTTTT | Amplification of pUC19 |
| *cph2*-LA-2 | ATCCCGCCGCCGCCGCCGGATGTTTTGACTCGGCGGGGATT | Amplification of *cph2*-LA |
| *cph2*-LA-F | AAAACGACGGCCAGTGAATTCGGGGGATGGATAGGAAGAGCAA | Amplification of *cph2*-LA |
| *cph2*-RA-2 | GACCATGATTACGCCAAGCTTGCGGTAACTAGCGGAGCGGC | Amplification of *cph2*-RA |
| *cph2*-RA-1 | ATCCGCGCGCGCGCGCGCGATCCCGATCGCCTAACAATTAAGCGG | Amplification of *cph2*-RA |
| P*aphI*-GC-R | ATCGCGCGCGCGCGCGCGGAT | Amplification of *aphI* |
| P*aphI*-C2G-F | ATCCGGCGGCGGCGGCGGGAT | Amplification of *aphI* |
| PpSCT2R6970 | CCAAAAAAAAACCCCGCCGA | Amplification of pSCT1(170831X2) |
| PpSCT2F7401 | TTCGGTGATACCAGCATCGT | Amplification of pSCT1(170831X2) |
| P*sll0199*R9m | ATGCTGGTATCACCGAAGCGATTGTATCTATAGGGACTTGT | Amplification of *Synechocystis* 6803 *petE* promoter for pSCT3 |
| P*sll0199*F438m | GCGGGGTTTTTTTTTGGTGCAAGGATTCATAGCGGTTG | Amplification of *Synechocystis* 6803 *petE* promoter for pSCT3 |
| PpSCT-*slr1970*F1 | GAGGTAACAACAAGATGAGCATCCAAGAAATCTTTACC | Amplification of *slr1970* |
| PpSCT-*slr1970*R519 | ACCACCAGAACCCCCATCCTGGGCTAGGGCT | Amplification of *slr1970* |
| PV_19 | CATCTTGTTGTTACCTCCTTAGCA | Linearization of pSCT3(180515X3) |
| PV_20 | GGGGGTTCTGGTGGTGGTAGCACT | Linearization of pSCT3(180515X3) |
| P*ydeH*F4 | AAGGAGGTAACAACAAGATGATCAAGAAGACAACGGAAAT | Amplification of *ydeH* |
| P*ydeH*R891-PV26 | ACCACCAGAACCCCCTTAAACTCGGTTAATCACATTTT | Amplification of *ydeH* |
| PV_26 | GGTGGATCTGGAGGTAGTGGTTCTGGTGGTGGTAGCAC | Amplification of pSCT3(180515X3) |
| P*yhjH*F4 | AAGGAGGTAACAACAAGATGATAAGGCAGGTTATCCAGCGAA | Amplification of *yhjH* |
| P*yhjH*R768-PV26 | ACCACCAGAACCCCCTTATAGCGCCAGAACCGCCGT | Amplification of *yhjH* |

**Supplementary Table S2.** Plasmids used in this study.

| **plasmid** | **description** | **reference** |
| --- | --- | --- |
| pUC19-cdgR-KmR | AmpR, KmR; used for deletion of the *cdgR/slr1970* | This study |
| pEP-cdgR-FLAG | GenR; expression of Hfq and CdrG-FLAG from its native promoter, used for complementation | This study |
| pCpf1b | SpR SmR; vector carrying the CRISPR-pCpf1c genome editing system and *Bacillus subtilis* *sacB* for counter selection | Niu et al. 2019 |
| pCpf1b-cdgR-sp | SpR SmR; used for markerless deletion of *cdgR/slr1970* | This study |
| pCpf1-Mslr1970R236 | SpR SmR; used for markerless deletion of *cdgR/slr1970*, without *sacB* | This study |
| pUC19-*cph2*-km | KmR NeoR; used for deletion of the *cph2* gene | This study |
| pCT | synthetic *petE* promoter from *Anabaena* sp. PCC 7120 coupled to a theophylline riboswitch, inducible by copper and theophylline | Xing et al., 2020 |
| pSCT3 | Modified pSCT plasmid, Anabaena P*_petE_* replaced by *Synechocystis* 6803 P*_petE_*, synthetic promoter, inducible by copper and theophylline | This study |
| pSCT3-*cdgR* | KmR, NeoR; expression of CdgR | This study |
| pSCT3-*ydeH* | KmR, NeoR; expression of *E. coli* YdeH | This study |
| pSCT3-*yhjH* | KmR, NeoR; expression of *E. coli* YhjH | This study  Zeng et al., 2023 |
| pJET-*∆crhR*-SpecR | AmpR, SpecR; inactivation of RNA helicase CrhR; used for determination of transformation efficiency | This study  Prakash et al.2010 |
| pKT25 | KmR; Two-hybrid plasmid, C-terminal CyaA (T25) fusion | Karimova et al, 1998 |
| pKNT25 | KmR; Two-hybrid plasmid, N-terminal CyaA (T25) fusion | Karimova et al, 1998 |
| pUT18 | AmpR; Two-hybrid plasmid, C-terminal CyaA (T18) fusion | Karimova et al, 1998 |
| pUT18C | AmpR; Two-hybrid plasmid, N-terminal CyaA (T18) fusion | Karimova et al, 1998 |
| pKT25-zip | KmR; pKT25 in which the leucine zipper of *yeast* GCN4 is genetically fused in-frame to the T25 fragment | Karimova et al, 1998 |
| pUT18C-zip | AmpR; pUT18C in which the leucine zipper of *yeast* GCN4 is genetically fused in-frame to the T18 fragment | Karimova et al, 1998 |
| pKT25-*cdgR* | KmR; pKT25 carrying CdgR fused in frame to T25 at the N-Terminus | This study |
| pKNT25-*cdgR* | KmR; pKT25 carrying CdgR fused in frame to T25 at the C-Terminus | This study |
| pUT18-*cdgR* | AmpR; pUT18 carrying CdgR fused in frame to T18 at the C-Terminus | This study |
| pUT18C-*cdgR* | AmpR; pUT18C carrying CdgR fused in frame to T18 at the N-Terminus | This study |
| pKT25-*sycrp1* | KmR; pKT25 carrying SyCRP1 fused in frame to T25 at the N-Terminus | This study |
| pKNT25-*sycrp1* | KmR; pKNT25 carrying SyCRP1 fused in frame to T25 at the C-Terminus | This study |
| pUT18-*sycrp1* | AmpR; pUT18 carrying SyCRP1 fused in frame to T18 at the C-Terminus | This study |
| pUT18C-*sycrp1* | AmpR; pUT18C carrying SyCRP1 fused in frame to T18 at the N-Terminus | This study |
| pKT25-*sycrp2* | KmR; pKT25 carrying SyCRP2 fused in frame to T25 at the N-Terminus | This study |
| pKNT25-*sycrp2* | KmR; pKNT25 carrying SyCRP2 fused in frame to T25 at the C-Terminus | This study |
| pUT18-*sycrp2* | AmpR; pUT18 carrying SyCRP2 fused in frame to T18 at the C-Terminus | This study |
| pUT18C-*sycrp2* | AmpR; pUT18C carrying SyCRP2 fused in frame to T18 at the N-Terminus | This study |
| pUT18-*bicA* | AmpR; pUT18 carrying BicA fused in frame to T18 at the C-Terminus | Haffner et al., 2025 |
| pKT25-*sbtB* | KmR; pKT25 carrying SbtB fused in frame to T25 at the N-Terminus | Selim et al., 2021 |
| pUT18C-*glgB* | AmpR; pUT18C carrying GlgB fused in frame to T18 at the N-Terminus | Selim et al., 2021 |
| pUT18-*glgC* | AmpR; pUT18 carrying GlgC fused in frame to T18 at the C-Terminus | Haffner et al., 2025 |

**Supplementary Table S3.** Strains used in this study.

| **Host** | **strain** | **purpose** | **reference** |
| --- | --- | --- | --- |
| *Synechocystis* sp. PCC 6803 | WT |  | Trautmann et al., 2012 |
|  | *cdgR::Km* | inactivation of *cdgR/slr1970* | This study |
|  | *∆cdgR* | markerless inactivation of *cdgR/slr1970* | This study |
|  | *cdgR::Km/ cdgR-FLAG* | complementation strain of *cdgR::Km*, expression of *cdgR*-FLAG from ist native promoter | This study |
|  | *WT_OE cdgR* | overexpression of *cdgR* | This study |
|  | WT_OE *yhjH* | overexpression of *E. coli yhjH* | This study |
|  | WT_OE *ydeH* | overexpression of *E. coli ydeH* | This study |
|  | Δ*cdgR*_OE *yhjH* | overexpression of *E. coli yhjH* in the *∆cdgR* background | This study |
|  | Δ*cdgR*_OE *ydeH* | overexpression of *E. coli ydeH* in the *∆cdgR* background | This study |
|  | *C-∆cdgR* | complementation strain of *∆cdgR*, expression of *cdgR*-FLAG from its native promoter | This study |
|  | *∆cph2/* ∆*cdgR* | inactivation of *cph2* in *∆cdgR/slr1970* | This study |
|  | *∆cph2* | inactivation of *cph2* | This study |
| *Escherichia coli* | BL21(DE3) | protein expression | Studier & Moffatt, 1986 |
|  | DH5α | cloning and plasmid propagation | Grant et al., 1990 |
|  | TOP10F' | cloning and plasmid propagation of BATCH related constructs | Invitrogen |
|  | J53/RP4 | *E. coli* helper strain for tri-parental conjugation | Datta et al., 1971 |
|  | BTH101 | *E. coli* strain lacking cAMP cyclase CyaA, used in BATCH assay | Battesti & Bouveret, 2012 |
